# Unraveling the complex story of intergenomic recombination in ABB allotriploid bananas

**DOI:** 10.1101/830059

**Authors:** Cenci Alberto, Sardos Julie, Hueber Yann, Martin Guillaume, Breton Catherine, Roux Nicolas, Swennen Rony, Carpentier Sebastien Christian, Rouard Mathieu

## Abstract

**Background and Aims:** Bananas (*Musa* spp.) are a major staple food for hundreds of millions of people in developing countries. The cultivated varieties are seedless and parthenocarpic clones of which the ancestral origin remains to be clarified. The most important cultivars are triploids with an AAA, AAB, or ABB genome constitution, with A and B genomes provided by *M. acuminata* and *M. balbisiana*, respectively. Previous studies suggested that inter-genome recombinations were relatively common in banana cultivars and that triploids were more likely to have passed through an intermediate hybrid. In this study, we investigated the chromosome structure within the ABB group, composed of starchy cooking bananas that play an important role in food security.

**Methods:** Using SNP markers called from RAD-Seq data, we studied the chromosome structure of 36 ABB genotypes spanning defined taxonomic subgroups. To complement our understanding, we search for similar events within nine AB hybrid genotypes.

**Key Results:** Recurrent Homoeologous Exchanges (HEs), i.e. chromatin exchanges between A and B subgenomes were unraveled with at least 9 founding events at the origin of the ABB bananas prior to the clonal diversification. The discovery of this nine founding events allows discussing the possible routes that led to the creation of the different subgroups and formulate new hypotheses. Based on our observations, we suggest different routes that gave rise to the current diversity in the ABB cultivars. Routes involving primary AB hybrids, routes leading to shared HEs and routes leading to a B excess ratio. Genetic fluxes took place between *M. acuminata* and *M. balbisiana*, particularly in India, where these unbalanced AB hybrids and ABB allotriploid originated and where cultivated *M. balbisiana* are abundant.

**Conclusions:** The result of this study clarifies the classification of ABB cultivars and leading possibly to the revision of the classification of this subgroup. This is an important step to unravel the origin of polyploid bananas, and contributes to possible scenarios on the origin. ABB bananas are hypothesized to be more drought tolerant. Knowing the origin of our current cultivars and so their potential parents will help breeders to make the right choices for future crosses. The *M. balbisiana* genome is a good source to create new cultivars able to answer the numerous challenges of banana breeding.

## Introduction

Bananas (*Musa* spp.) are herbaceous monocotyledons belonging to the Zingiberales. The *Musa* genus originated in south-east Asia and west Oceania where it was domesticated; from there banana was spread to tropical areas of Africa and America. Two species widely contributed to the varieties cultivated worldwide: *Musa acuminata* Colla (A genome), whose distribution coincides with Mainland and Maritime Southeast Asia and New-Guinea island and *M. balbisiana* Colla (B genome), distributed in the eastern South Asia, northern Southeast Asia, and southern China (Simmonds 1962; Janssens *et al.* 2016).

Present day cultivated bananas with exclusively A genome constitution are noted as AA or AAA and hybrids noted as AB, AAB or ABB, depending to their ploidy level. AA diploids are parthenocarpic with various degrees of sterility, mainly cultivated for fruit consumption. Edible AB hybrids do exist but are rare, mostly known in India. Those studied so far are reported as sterile (Jenny *et al.* 2011). *M. balbisiana*, although associated with wild populations, can be found in home gardens where it is cultivated for its leaves, male bud, young fruits or seeds, the latest being used in traditional pharmacopoeia (Subbaraya *et al.* 2006). The most common banana cultivars are triploid (2n=3x=33), i.e. composed of three sets of 11 chromosomes and are either autotriploid (AAA) or allotriploid (AAB and ABB). These cultivars have very low seed set due to farmers’ selection combined with triploidy that hampers the production of balanced gametes. Consequently, they are clonally propagated. The combination of sterility and parthenocarpy ensures the production of seedless fruits that are edible.

The taxonomic classification of allotriploid bananas was mainly based on morphological descriptors with traits differentiating *M. acuminata* from *M. balbisiana* (Simmonds and Shepherd 1955). Later, molecular analysis of organelle genomes allowed the characterization of cultivar cytotypes and, combined with ploidy measurements, lead to hypothesis on crossing pathways (Carreel *et al.* 2002; Boonruangrod *et al.* 2008). Next molecular markers from Single sequence repeats (SSRs) provided a multi-locus survey of the parental allele contribution (Hippolyte *et al.* 2012; Christelová *et al.* 2017). Cytogenetics and genome-wide studies also revealed that allotriploids are not mere additions of sets of 11 chromosomes, but contain regions with a variable subgenome ratio along the chromosomes (D’Hont *et al.* 2000; Jeridi *et al.* 2011; Noumbissié *et al.* 2016; Baurens *et al.* 2019; Cenci *et al.* 2019), demonstrating the occurrence of recombinations between subgenomes. It is commonly accepted that triploids have been generated by spontaneous hybridizations of edible AA or AB with various sources of the additional A or B-genome (other edible AA or *M. acuminata*, and *M. balbisiana*). However, the above cited studies point out the genomic complexity of allotriploid cultivars, suggesting the occurrence of backcrosses with parental species through residual fertility in some allotriploid cultivars (De Langhe *et al.* 2010).

Among the allotriploids, the ABB genomic group comprises starchy bananas that are used for cooking, dessert and beer production (Karamura *et al.* 1998). They are reported of being resistant to weevil, nematodes, Black leaf streak (Karamura *et al.* 1998) and tolerant to drought (Thomas *et al.* 1998; De Langhe 2002; Vanhove *et al.* 2012; Ravi *et al.* 2013; Kissel *et al.* 2015; van Wesemael *et al.* 2019). The ABB banana cultivars belong to a morphologically diverse group subdivided into 9 subgroups namely Bluggoe, Monthan, Ney Mannan, Klue Teparod, Kalapua, Peyan, Pisang Awak, Pelipita and Saba (Daniells *et al.* 2001). They originate from two regions: India and surrounding regions, and South East Asia (De Langhe *et al.* 2009; Perrier *et al.* 2011). Molecular markers have shown that some ABB cultivars were erratically classified (Sardos *et al.* 2016; Christelová *et al.* 2017). Furthermore, the genomic composition of the Saba subgroup, coming from the Philippines, has been debated for years as some authors suggested it consists of B genomes only (Valmayor *et al.* 1999).

The goal of this study was i) to survey a large sample of interspecific AB and ABB genotypes to establish molecular karyotypes and thus to set up a clear framework for their classification and ii) to bring new insights into origin and evolution of ABB triploid cultivars.

## Materials and Methods

### Plant material

Lyophilized leaf samples from 45 banana genotypes (36 allotriploid ABB and 9 hybrids AB) were supplied by the Bioversity International *Musa* germplasm Transit Centre (ITC) hosted at KU Leuven, Belgium except for three leaf samples collected in Indonesia during the Triangle collecting missions in 2012 – 2013 (Hermanto *et al.* 2014; Hermanto *et al.* 2014) originally considered to diploid AA accessions but revised as diploid AB (Christelová *et al.* 2017) and currently conserved at the Indonesian Centre for Horticultural Research and Development (ICHORD). The passport data of these accessions are available in the Musa Germplasm Information System (MGIS) (Ruas *et al.* 2017) (**Table 1**).

**Table 1:** List of 36 ABB and 9 AB accessions used in this study. Cytotypes: Ca and Cb indicate chloroplasts originating from *M. acuminata* and *M. balbisiana*, respectively; Ma and Mb indicated the same for mitochondria.

| Accession code | Accession name | DOI | Genome | Subgroup | Collection | Geographical origin | Cytotype | Genomic pattern |
| --- | --- | --- | --- | --- | --- | --- | --- | --- |
| ITC0245 | Safet Velchi | 10.18730/9 JM13 | AB | Ney Poovan | ITC | India | CaMa | 2x-1 |
| ITC1034 | Kunnan | 10.18730/9 M2KD | AB | Ney Poovan | ITC | India<sup>§</sup> | CaMa | 2x-2 |
| ITC1747 | Agniswar | 10.18730/9 NGN7 | AB | Kunnan | ITC | Indonesia | - | 2x-2 |
| ITC1729 | Padali Moongil | 10.18730/9 NFE5 | AB | Kunnan | ITC | India<sup>§</sup> | - | 2x-3 |
| ITC1752 | Poovilla Chundan | 10.18730/9 NH1K | AB | Kunnan | ITC | India<sup>§</sup> | - | 2x-3 |
| ITC1751 | Adukka Kunnan | 10.18730/9 NGYG | AB | Kunnan | ITC | India<sup>§</sup> | - | 2x-4 |
| Sum002 | Muku Bugis | - | AB | - | ICHORD | Indonesia | - | (2x) |
| Sum004 | Mu'u Seribu | - | AB | - | ICHORD | Indonesia | - | (2x) |
| ITC1880 | Mu'u Pundi | 10.18730/P 5G84 | AB |  | ITC | Indonesia (Sup008) | - | (2x) |
| ITC1700 | Kepok Kuning | 10.18730/9 ND89 | ABB | Saba | ITC | Indonesia | - | N/A |
| ITC1745 | Kepok Tanjung | 10.18730/9 NGH3 | ABB | Saba | ITC | Indonesia | - | N/A |
| ITC0632 | Cachaco enano | 10.18730/9 KATX | ABB | Bluggoe | ITC | Colombia* | - | 1a |
| ITC0643 | Cachaco | 10.18730/9 KBJG | ABB | Bluggoe | ITC | Colombia* | - | 1a |
| ITC1728 | Sambrani Monthan | 10.18730/9 NFD4 | ABB | Monthan | ITC | India | - | 1a |
| ITC1746 | Bankel | 10.18730/9 NGM6 | ABB | Pisang Awak | ITC | India | - | 1a |
| ITC1748 | Boddida Bukkisa | 10.18730/9 NGQ9 | ABB | Pisang Awak | ITC | India | - | 1a |
| ITC1483 | Monthan | 10.18730/9 MXQY | ABB | Monthan | ITC | India | CaMb | 1a |
| ITC0026 | Sabra | 10.18730/9 J6SQ | ABB | Unknown | ITC | Gabon | - | 1a+ |
| ITC0767 | Dole | 10.18730/9 | ABB | Bluggoe | ITC | France* | CaMb | 1a+ |
| | | KGR\$ | | | | | | |
| ITC0361 | Blue Java | 10.18730/9<br>JTR~ | ABB | Ney<br>Mannan | ITC | Fiji* | - | 1b |
| ITC1750 | Ney Vannan | 10.18730/9<br>NGXF | ABB | Ney<br>Mannan | ITC | India | - | 1b |
| ITC1738 | Kyauk Sein<br>Phee Kyan | 10.18730/9<br>NG4V | ABB | Unknown | ITC | Myanmar | - | 1b |
| ITC0123 | Simili Radjah | 10.18730/9<br>JC16 | ABB | Peyan | ITC | India | CbMb | 1b+ |
| ITC1600 | INIVIT PB-2003 | 10.18730/9<br>N676 | ABB | Saba | ITC | Cuba* | - | 1c |
| ITC1138 | Saba | 10.18730/9<br>M6MZ | ABB | Saba | ITC | Philippines | CaMb | 1c |
| ITC0659 | Namwa Khom | 10.18730/9<br>KCB4 | ABB | Pisang<br>Awak | ITC | Thailand | CbMb | 2 |
| ITC0053 | Bom | 10.18730/9<br>J7YQ | ABB | Pisang<br>Awak | ITC | Ivory<br>Coast* | - | 2 |
| ITC0087 | Kayinja | 10.18730/9<br>J9YD | ABB | Pisang<br>Awak | ITC | Burundi* | - | 2 |
| ITC0101 | Fougamou 1 | 10.18730/9<br>JAX7 | ABB | Pisang<br>Awak | ITC | Gabon | - | 2 |
| ITC0526 | Kluai Namwa<br>Khom | 10.18730/9<br>K3PQ | ABB | Pisang<br>Awak | ITC | Thailand* | - | 2 |
| ITC1599 | Kambani<br>Zambia | 10.18730/9<br>N665 | ABB | Pisang<br>Awak | ITC | Tanzania | - | 2 |
| ITC1721 | Karpuravalli | 10.18730/9<br>NEWR | ABB | Pisang<br>Awak | ITC | India | - | 2 |
| ITC1737 | Ya Khine | 10.18730/9<br>NG1R | ABB | Unknown | ITC | Myanmar | - | 2 |
| ITC0339 | Pisang Awak | 10.18730/9<br>JSGY | ABB | Pisang<br>Awak | ITC | Australia* | - | 2+ |
| ITC1719 | Chinia | 10.18730/9<br>NERM | ABB | Pisang<br>Awak | ITC | India | - | 3 |
| ITC1749 | Vananthpurani | 10.18730/9<br>NGTC | ABB | Pisang<br>Awak | ITC | India | - | 3 |
| ITC0472 | Pelipita | 10.18730/9<br>K0MU | ABB | Unknown | ITC | Philippines | CaMb | 4 |
| ITC0396 | Pelipita | 10.18730/9<br>JW32 | ABB | Unknown | ITC | Costa Rica* | - | 4 |
| ITC0397 | Pelipita<br>Majoncho | 10.18730/9<br>JW65 | ABB | Unknown | ITC | Costa Rica* | - | 4 |
| ITC0652 | Kluai Tiparot | 10.18730/9 | ABB | Unknown | ITC | Thailand | CbMb | 5 |
|  |  | KC0Y |  |  |  |  |  |  |
| ITC0473 | Balonkawe | 10.18730/9<br>K0P1 | ABB | Klue<br>Teparod | ITC | Philippines | - | 5 |
| ITC0983 | Auko | 10.18730/9<br>KZKW | ABB (M) | Unknown | ITC | Papua N. G. | - | 6 |
| ITC0987 | Auko | 10.18730/9<br>KZZ3 | ABB (M) | Unknown | ITC | Papua N. G. | - | 6 |
| ITC0990 | Vunapope | 10.18730/9<br>M06A | ABB (M) | Unknown | ITC | Papua N. G. | - | 6 |
| ITC1682 | Chuoi mit | 10.18730/9<br>NC06 | ABB<br>(AB) | Unknown | ITC | Vietnam | - | 7 |

\*Geographical origin of the previous *ex situ* collection; +Genomic pattern containing aneuploidy regions;
§ Assumed geographical origin from literature or name; (AB) Genome erroneously classed or labelled as AB in Taxonomic classification (passport data) but recognized as triploid in Christelova *et al.* (2017); (M) erroneously classed as AB in Taxonomic classification and reported as mixoploidy (2x+3x); + Pattern containing accession specific aneuploidy.

### DNA extraction and RAD-Seq data generation

Genomic DNA was extracted using 2X CTAB protocol. The method used to create the library for restriction-site associated DNA sequencing (RAD-Seq) used the *Pst*I restriction enzyme. The 300-500 short-insert libraries were sequenced with 91 bp pair-end reads using Illumina HiSeq2000 (Illumina, San Diego, CA, USA) by BGI Hong Kong. At BGI, the raw data were modified with the following two steps: (1) reads polluted by adapter sequences were deleted and reads that contained >50% low-quality bases (quality value ≤ 5) or >10% N bases were removed.

### Read mapping, filtering and SNP calling

Pair-end reads contained in raw FASTQ files were checked using FastQC. Reads were then cleaned to remove adapter sequences and low-quality ends (Phred score > 20) with Cutadapt (Martin 2011). After trimming, reads inferior to 30 bp were discarded. Reads were then aligned using BWA (Li and Durbin 2009) with default parameter against the *Musa acuminata* genome of the reference v2 (DH Pahang) (D’Hont *et al.* 2012; Martin *et al.* 2016) downloaded on the Banana Genome Hub (Droc *et al.* 2013). Read groups were added for each alignment and reads were locally realigned with IndelRealigner (Genome Analysis ToolKit) (McKenna *et al.* 2010). HaplotypeCaller from GATK version 3.4-46 was then used to get a gVCF file for each accession. The genotyping on gVCF files were performed with GenotypeGVCFs from GATK 3.4-46. SNPs were called on uniquely mapped reads with HaplotyCaller from GATK version 3.4-46.

### Detection of Homoeologous Exchange (HE)

The subgenomic structure was investigated by VcfHunter (https://github.com/SouthGreenPlatform/VcfHunter). Based on known sequence variability in the A and B genomes, SNP variants were assigned to the ancestral genomes in order to plot the A/B genome allele coverage ratio and to calculate the normalized site coverage along chromosomes as described in Baurens *et al.* 2019. SNP datasets from accessions representing *M. acuminata* and *M. balbisiana* sequence variability were retrieved with GIGWA (Sempéré *et al.* 2016) on MGIS (https://www.crop-diversity.org/mgis/gigwa) (Ruas *et al.* 2017). HEs were inferred when, in a given chromosome, a change in A/B allele ratio was observed between adjacent regions. Chromosomes were identified by their number in the A genome reference (from 01 to 11) (D’Hont *et al.* 2012) proceeded by “chr”. Chromosome arms were defined as first or second according to the chromosome sequence.

## Results

### Molecular karyotyping

A set of 9 AB and 36 ABB genotypes were used to identify SNPs markers from RAD-Seq datasets. The number of SNPs ranged between 10,000 and 26,000 per chromosome for each genotype (182,000 SNPs by genotype in average) (**Supplementary Material 1**). Similarly as Baurens *et al.* (2019), SNPs were assigned to the A and B subgenomes and used to scan chromosomes for the detection of deviating regions, i.e. portions of chromosomes where the subgenome ratios differ from the expected based on the assumed genomic composition (i.e. A1:B1 and A1:B2 for the hybrids AB and the allotriploid ABB, respectively) also referred as homologous exchanges (HEs).

#### AB hybrids

A panel of 9 banana accessions reported as AB hybrids was analyzed (**Table 1**). Only one third had the expected subgenome composition (i.e. A1:B1), while the others (6) showed one or two deviating regions with A2:B0 ratio (i.e. missing B-specific SNPs). By comparing all HE together, we delineated four patterns denoted as 2x-1, 2x-2, 2x-3 and 2x-4. The cultivar ‘Safet Velchi’ had a A2:B0 ratio in the first arm terminal region of chr06 (**Figure 1**, pattern 2x-1); cultivars ‘Kunnan’ and ‘Agniswar’ showed A2:B0 ratio in the terminal region of second arm of chr08 and in a first arm interstitial region of chr09 (**Figure 1**, pattern 2x-2). ‘Padali Moongil’ and ‘Poovilla Chundan’ showed A2:B0 ratio in the first arm terminal region of chr04 and in a first arm interstitial region of chr09 (**Figure 1**, pattern 2x-3); ‘Adukka Kunnan’ had an interstitial A2:B0 region in chr09 first arm (**Figure 1**, pattern 2x-4). Among those samples, the cytotypes of ‘Safet Velchi’ and ‘Kunnan’ were previously characterized with chloroplast and mitochondrion originating from *M. acuminata* (Carreel *et al.* 2002; Boonruangrod *et al.* 2008).

**Figure 1.**
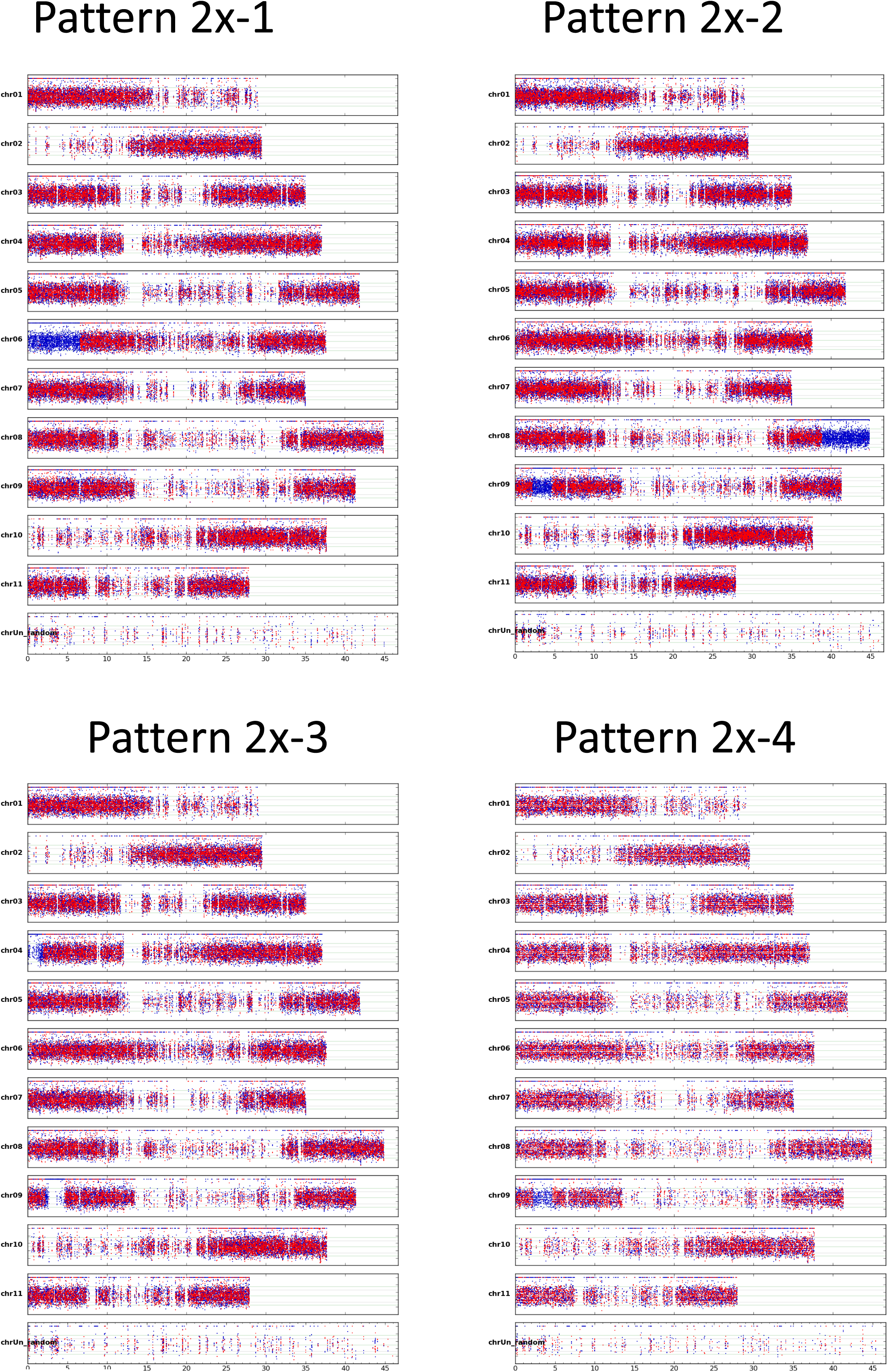
Patterns of AB hybrids having genomic regions with unexpected subgenome ratio. Coverage ratio of SNP (y axis) assigned to A and B genomes are marked in blue and red, respectively. SNP coordinates (x axis) are reported in Mb at the bottom of each genotype figure.

#### ABB allotriploid

In 34 out of the 36 ABB genotypes, HEs with subgenome ratio corresponding to A0:B3 (region with only B assigned SNPs) or A2:B1 (region with 2/3 A -1/3 B assigned SNPs) were detected. None with A3:B0 ratio were found. Nine patterns corresponding to specific combinations were identified, enabling the classification of each genotype (**Table 1**). Three patterns sharing regions with the same ratio deviation and size were numbered 1a, 1b and 1c. In those, large terminal regions of the second arm of chr04 and chr11 have an A0:B3 ratio and the first arm of chr09 shows an A2:B1 interstitial region (**Figure 2**). Patterns 1a and 1c have in common the A0:B3 terminal region of the chr11 first arm (**Figure 2**) but Pattern 1a differs with a specific interstitial region A2:B1 in chr04 first arm. Pattern 1c has three specific regions A0:B3 in chr02 second arm, chr04 first arm and in chr09 first arm. The interstitial region in chr09 where the A genome replaced the B counterpart has very a similar or identical size as the one shared by the 2x-2, 2x-3 and 2x-4 patterns in AB hybrids (**Figure 3**). These patterns contain mostly genotypes belonging to the Bluggoe, Monthan, Saba, Peyan and Ney Mannan subgroups that originated from India. The remaining patterns do not share identical HE and were named from 2 to 7. The pattern 2 and 3 gathers genotypes classified as Pisang Awak (**Figure 4**), with the exception of 2 assigned to the pattern 1a which are known to be erroneously allocated to this subgroup (Christelová *et al.* 2017). The patterns 4 and 5 are the only ones with entire chromosome substitution with the presence of an A0:B3 subgenome ratio. Specific to pattern 5, no regions with A2:B1 ratio were detected. Overall, the patterns 4 and 5 have an excess of B subgenome compared to other ones (**Figure 4**) and correspond to genotypes assigned to subgroups Pelipita and Klue Teparod, respectively. The pattern 6 is not assigned to any known subgroup but the 3 genotypes composing it share all the same origin from Papua New Guinea **(Figure 4, Table 1**). The pattern 7 is represented by only one genotype, ‘Chuoi mit’, coming from Vietnam **(Figure 4)**. Finally, two genotypes (‘Kepok Kuning’ and ‘Kepok Tanjung’, originally classified as belonging to subgroup Saba) both originating from Indonesia, do not show any region with deviating subgenome ratio and were not associated with any pattern.

**Figure 2.**
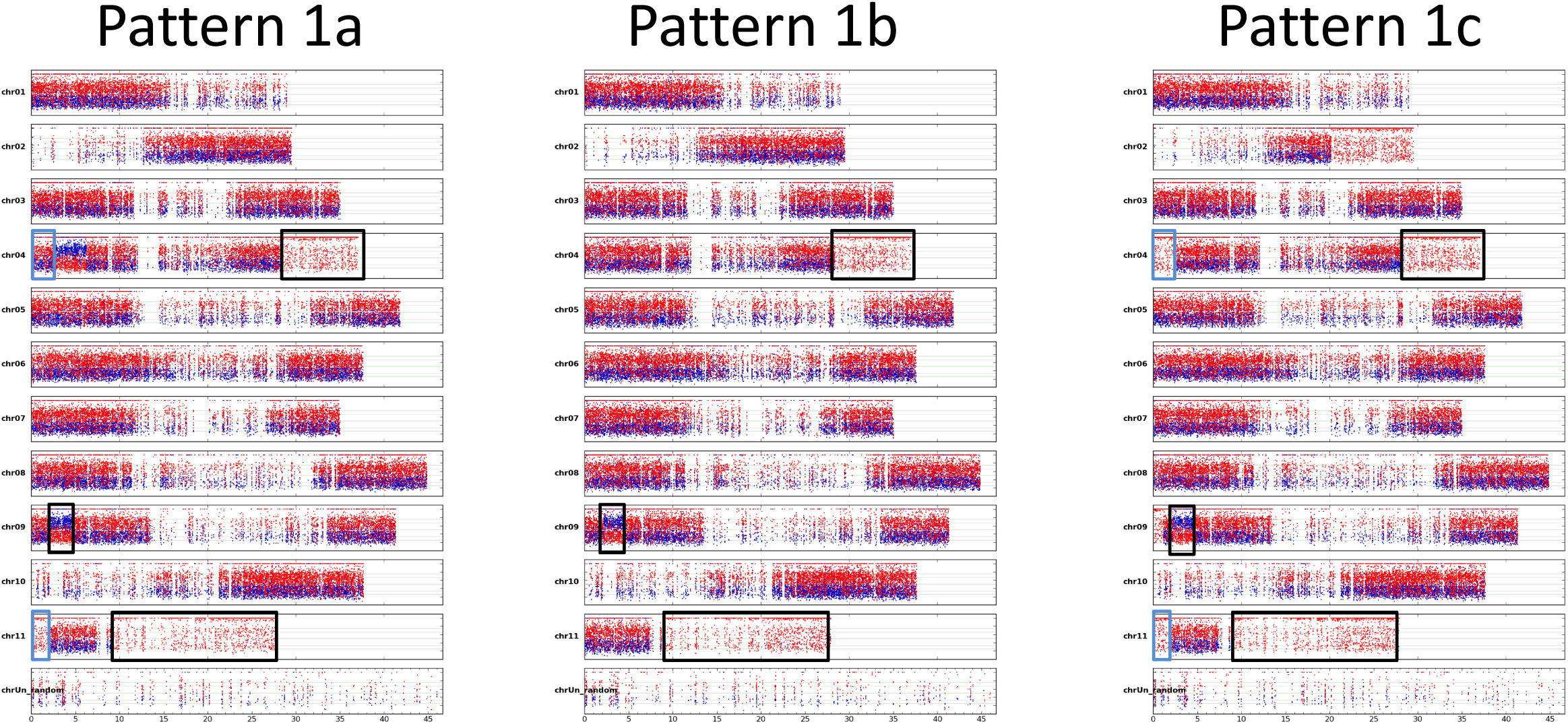
Patterns 1a, 1b and 1c of allotriploid ABB having genomic regions with unexpected allele ratio composition. Coverage ratio of SNP (y axis) assigned to A and B genomes are marked in blue and red, respectively. SNP coordinates (x axis) are reported in Mb at the bottom of each genotype figure. Black rectangles indicate regions shared by all three patterns, blue rectangles indicate regions shared by patterns 1a and 1c.

**Figure 3.**
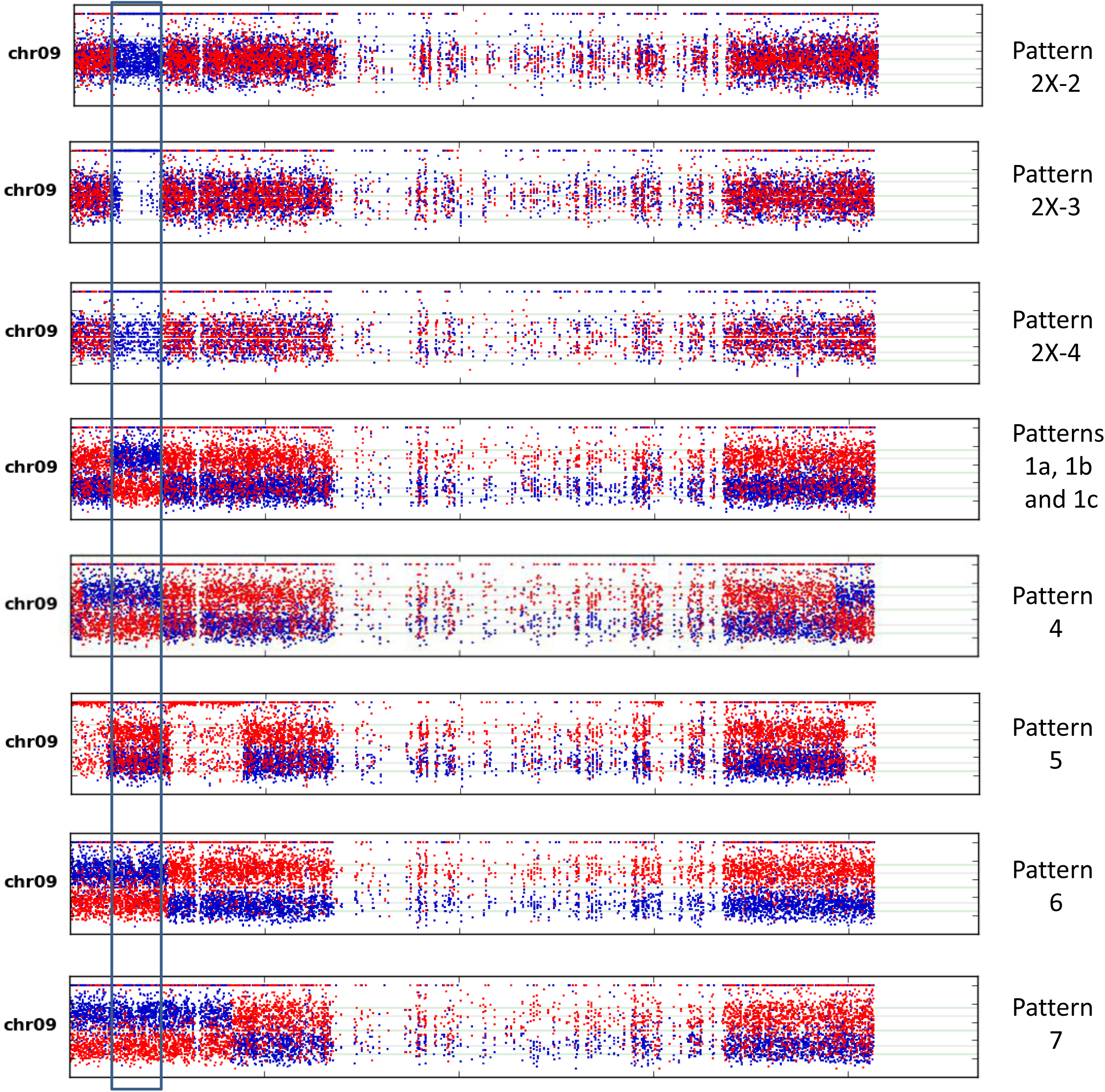
Chr09 of three AB and seven ABB patterns. The rectangle highlights the A genome enriched region.

**Figure 4.**
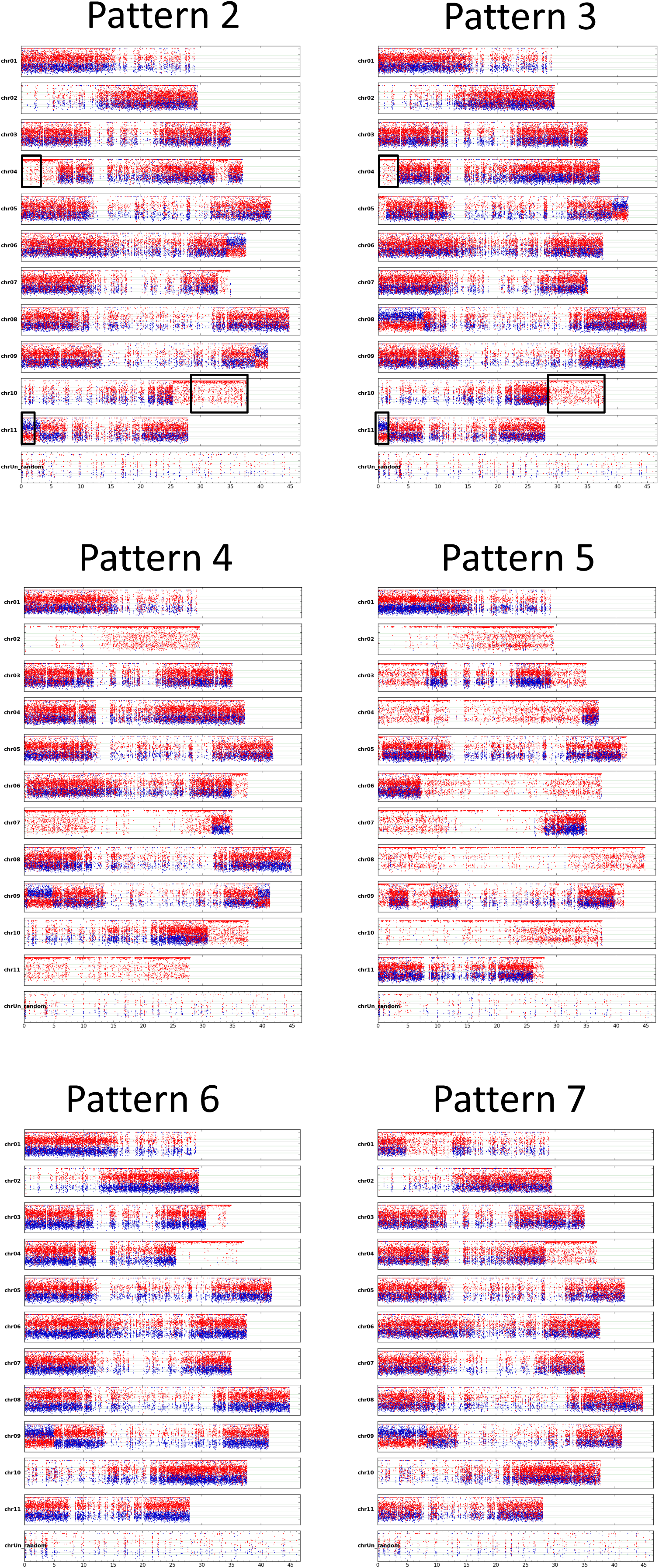
Patterns 2 to 7 of allotriploid ABB having genomic regions with unexpected allele ratio composition. Coverage ratio of SNP (y axis) assigned to A and B genomes are marked in blue and red, respectively. SNP coordinates (x axis) are reported in Mb at the bottom of each genotype figure. Black rectangles indicated region partially overlapping in the two patterns.

#### Genotype-specific aneuploidies

For 5 genotypes, in addition of the described HEs based on ratios involving three chromosomes (**Figure 5a**), portions or entire chromosomes displayed ratios with two or four copies. For the ‘Dole’ genotype belonging to the Bluggoe subgroup (pattern 1a), no SNPs assigned to the A subgenome were detected along the whole chr08. The lower SNP coverage compared to other chromosomes and the unimodal coverage distribution of residual B genome heterozygosity (at 0.5) indicates an A0:B2 ratio, implying the A version of chr08 was lost (**Figure 5b**). The chr06 of ‘Sabra’ (pattern 1a) shows a complex structure (**Figure 5c**). On the first arm, there is a similar coverage of A and B assigned SNPs around 0.5 and lower SNP coverage than other regions, compatible with an A1:B1 subgenome ratio. On the second arm, six regions with A1:B1, A1:B2 and A1:B3 ratios were inferred. In ‘Simili Radjah’, a large part of its second arm of the chr05 is missing. Most of the region exhibits a diploid pattern A1:B1 except in its terminal region where it appears A0:B2 (**Figure 5d**). The genotype ‘INIVIT PB-2003’ (pattern 1c) showed a unique interstitial region, in the chr10 second arm, where the SNPs assigned to A and B have similar contributions. Since SNP coverage in this region is higher than in the rest of chr10, an A2:B2 homeoallele ratio was inferred (**Figure 5e**). Finally, the accession ‘Pisang Awak’ (ITC0339, pattern 2) had an A1:B1 homeoallele ratio in the terminal part of the chr07 first arm (**Figure 5f**). Since these aneuploidies were specific to single genotypes (not shared with other genotypes with the same HE pattern), we considered that mutation events occurred after the original triploidization events.

**Figure 5.**
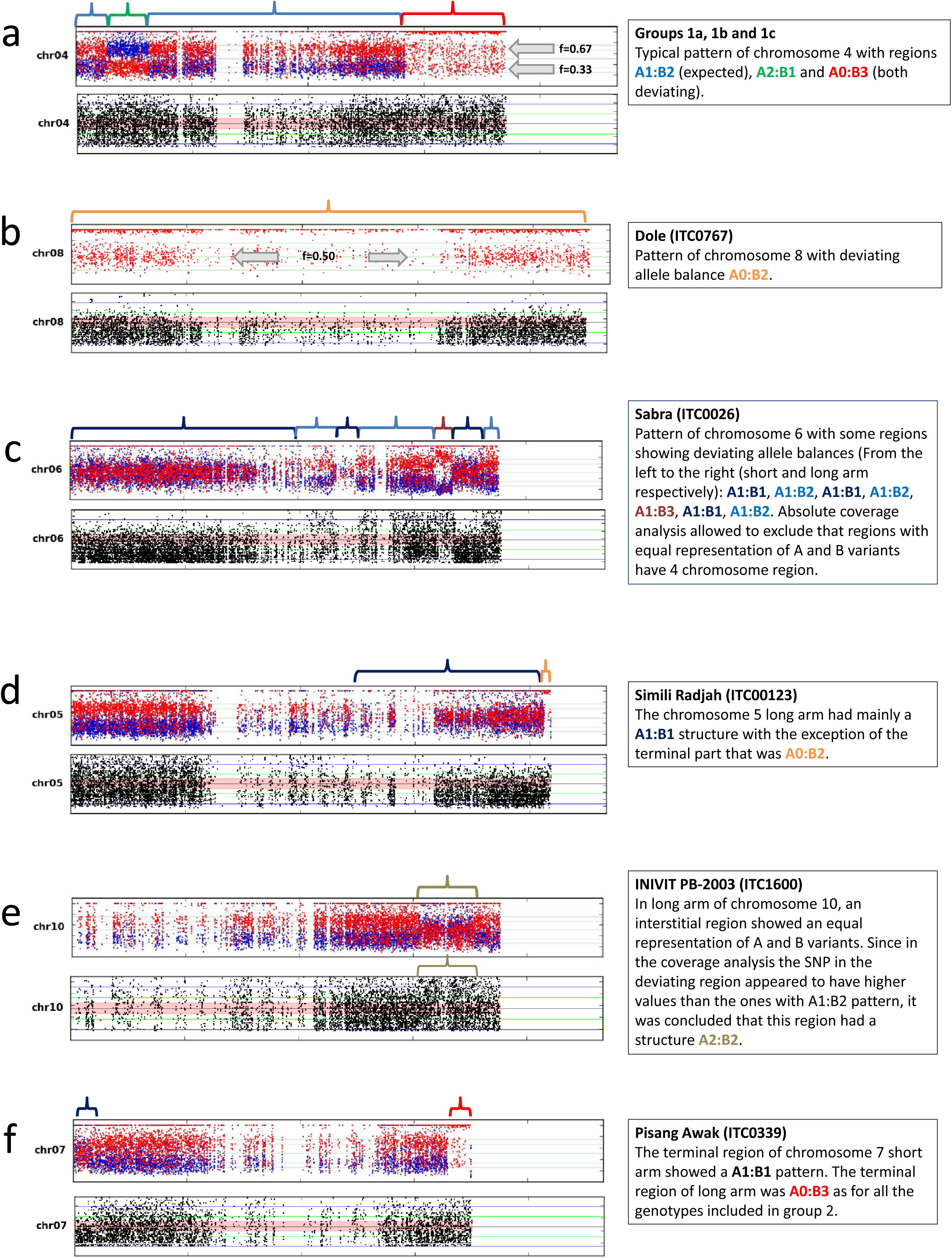
Pattern of critical chromosomes involved in aneuploidy. For each pattern, coverage ratio of SNP assigned to A and B subgenomes (in green and red, respectively) and absolute coverage are reported for chromosomes with deviating pattern. a) Pattern of chromosome present in expected number of copies (3); b-f) patterns with changes in number of copies involving entire or partial chromosomes. Curly bracket colors indicate regions with the inferred subgenome composition. Colored curly brackets delimitate the regions with inferred genome ratios reported in the right-side legend. Gray arrows illustrate different alleles frequencies used to discriminate between regions with A0:B3 and A0:B2 ratios in a and b patterns.

## Discussion

### The complex history of ABB cultivars can be grouped in at least nine founding events

The classification of cultivated ABB needs clarification since long. De Langhe 2002 noted that “The more popular names (Saba, Pisang Awak, Peyan, Bluggoe, Monthan) actually represent a cluster of closely related cultivars, generated by somatic variation. Each variant has its local name in Asia which makes nomenclature of the whole ABB group difficult to resolve based on solely on this source of information. The combination of morpho-taxonomic descriptors and SSR markers confirmed this difficulty to discriminate the Indian subgroups (Saraswathi *et al.* 2011), and the use of DArT or SSR markers on a wider sample showed that the classification was consistent for accessions belonging to the subgroup Pelipita, Klue Teparod and Pisang Awak (Sardos *et al.* 2016; Christelová *et al.* 2017). However, accessions classified belonging to the subgroups Saba, Monthan, Bluggoe, Ney Mannan or Peyan were often misclassified (Sardos *et al.* 2016).

In this study, genome-wide molecular data (RAD-Seq derived SNPs) were used to characterize the genome of diploid and triploid banana hybrid cultivars with emphasis on the ABB group. Overall, all the allotriploid accessions were confirmed having an ABB genome constitution, i.e. the ratio between variants assigned to B and A subgenomes was A1:B2. We further explored the presence of homoeologous exchanges (HEs) in a comprehensive set of ABB cultivars as initiated in previous studies (Baurens *et al.* 2019; Cenci *et al.* 2019) and confirmed that such a phenomenon has been very common in the creation of those domesticated bananas. Among the 36 genotypes surveyed, 34 had at least two chromosome portions for which the observed ratio was either A2:B1 or A0:B3 (**Table 2**; **Figure 6**).

**Table 2.**
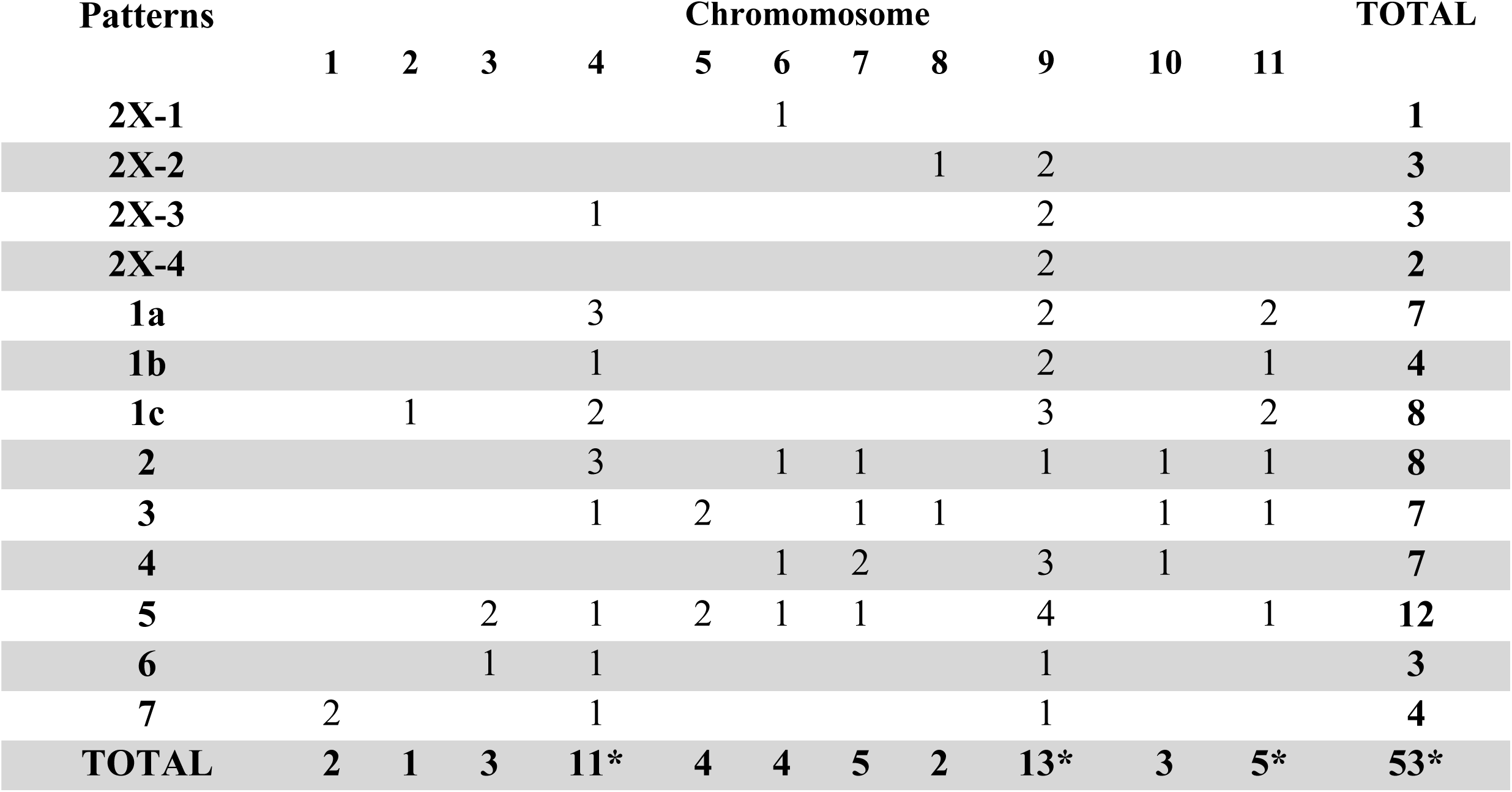
Number of Homoeologous Exchanges (HE) in ABB and AB patterns. HE are inferred when in a given chromosome a change in a subgenome ratio is observed (whole chromosome replacement are not counted). *Shared HEs are counted once.

**Figure 6.**
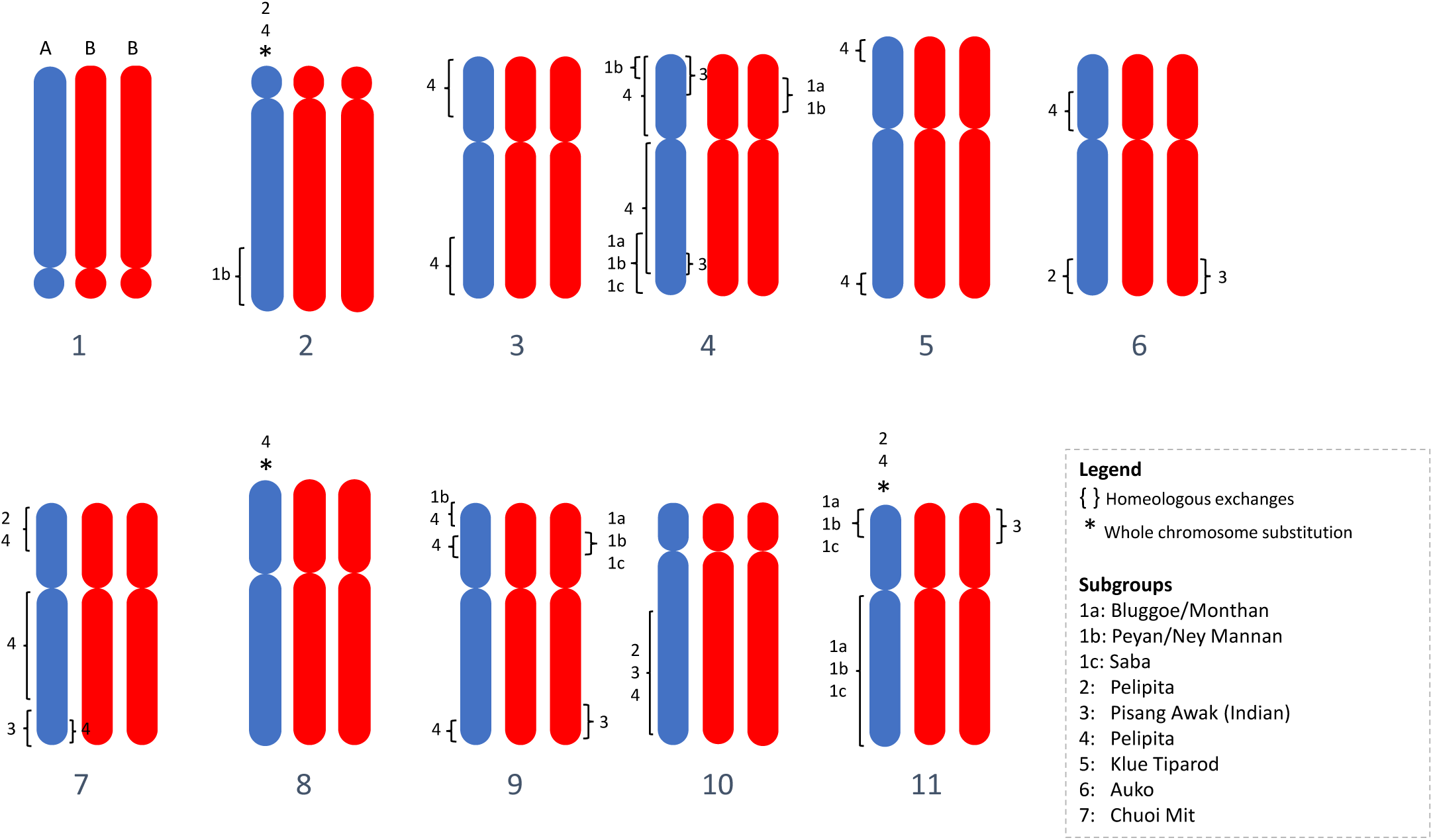
Ideogram overview of all HR and substitutions detected in 36 ABB bananas.

In total, 53 HEs were inferred in the 9 ABB and 4 AB patterns (shared HEs were counted only once) (**Table 2**). They were found in all eleven *Musa* chromosomes but are unevenly distributed, with 11 and 13 independent recombination events in the most recombined chr04 and chr09, respectively and only one recombination event in chr02 (**Table 2, Figure 6**). Since no recombination hotspots were observed for chr04 and chr09 by Baurens *et al.* (2019) using segregating populations, the high frequency of HE we detected in these chromosomes may be due to the presence of genes influencing domestication traits for which human selection could have favored some combinations between A and B homeoalleles. In particular, the first arm of chr09 appears to be involved in several independent HEs, and a subgenome A interstitial region was present in all the cultivars analyzed (**Figure 3**), suggesting the presence of an A homeoallele selected during the domestication process.

The different HE patterns of ABB allotriploidy provide new insights to their current classification:

### West ABB

#### Bluggoe, Monthan, Peyan and Ney Mannan share the same common origin

The dominance of accessions from India in patterns 1a and 1b is obvious. Equally, diploid hybrids (AB) from India appear to share HEs with patterns 1. We can therefore locate the early origin of the recombination events typical to patterns 1 in India. All cultivars of subgroup Bluggoe (**Table 1**), but also Monthan show the HE pattern 1a. Subgroups Peyan and Ney Mannan both have pattern 1b. This study shows that Bluggoe, Monthan, Peyan and Ney Mannan share a same common origin linked to India. Two additional Indian cultivars, classified as Pisang Awak, share the Pattern 3.

### East ABB

#### Pisang Awak subgroup would originate from at least two independent events

Eight Pisang Awak cultivars are comprised in pattern 2 (**Figure 4**). Comparing patterns 2 and 3 (i.e. coordinates of break points), we noticed that despite the independent occurrence of the recombination events in each pattern, similar deviations in A and B ratios are present in three chromosomal regions (black squares on **Figure 4**). We hypothesize that these regions harbor genes giving similar phenotypic traits to the cultivars with patterns 2 and 3 and that these traits influenced their classification in the same subgroup. In addition, phenotypic proximity among the cultivars of these two HE patterns likely results from genetic similarity in parental genotypes (Christelová *et al.* 2017).

#### Pelipita

All accessions (3) classified in the Pelipita subgroup shared an identical and specific deviating pattern (Pattern 4, **Figure 4**). These cultivars (‘Pelipita’ (ITC0396 and ITC0472) and ‘Pelipita Majoncho’), have entire A chr02 and chr11 substitution (A0:B3 structure) and the chr07 is mainly A0:B3 with the exception of the second arm interstitial region. These findings in ‘Pelipita’ genome are consistent with observations made through *in situ* hybridization reporting that ‘Pelipita’ contained 25 B and 8 A chromosomes (D’Hont *et al.* 2000). Additional chromosome regions with unexpected ratio are also present, notably two regions of chr09 exhibiting ratios favoring A (A2:B1).

#### Klue Teparod

The two cultivars of HE pattern 5 (‘Kluai Tiparot’ and ‘Balonkawe’) can be easily distinguished from all the other surveyed accessions. First, all deviating regions were A0:B3 (no A2:B1 were observed contrary to all other patterns), second, three entire chromosomes (chr02, chr08 and chr10) were being fully B and three additional A0:B3 centromere were present in recombined chromosomes (**Figure 4**). This observation provides new insights to a long-time debate that exists on the classification of the Klue Teparod subgroup. The ‘Balonkawe’ accession originally collected in the Philippines by Allen in 1959, was first classified as a tetraploid ABBB due to its robust appearance (Rosales *et al.* 1999). However, flow cytometry coupled with chromosomes counting assessed that the ‘Klue Tiparot’ accession from the same subgroup was in fact a triploid (Jenny *et al.* 1997). Later, the use of several types of molecular markers led to a different conclusion. Ribosomal DNA (Boonruangrod *et al.* 2008) suggested the presence of B genomes only while ITS sequences confirmed the occurrence of A (Hřibová *et al.* 2011; Čížková *et al.* 2013). Dominant DArT markers which resulted in the clustering of these two accessions into the *M. balbisiana* genepool led to hypothesize the presence of an incomplete A genome (Sardos *et al.* 2016). In this study, it is shown that Klue Teparod’s accessions contain a small fraction of an A genome, which makes this subgroup the cultivated variety carrying the highest content of B genome ever described. Like for the cultivar ‘Lep Chang Kut’ (ITC0647) registered as BBB in MGIS (Ruas *et al.* 2017) but for which the presence of A isoforms was detected (Carpentier *et al.* 2011), we provide evidence that Klue Teparod’s accessions are not parthenocarpic BBBs.

#### Saba

The four cultivars classified as Saba in our sample showed two different patterns: two Indonesian cultivars show a strict A1:B2 ratio whereas the other two of unknown origins show the 1c pattern. Although a bunch of cooking cultivars from the Philippines, including ‘Saba’ (not in our samples), were reported to be BBB (Valmayor *et al.* 1991; Sales *et al.* 2010), no such pattern was identified in the present study. However, it is confirmed that the same vernacular name is used for genotypes with different ancestral origins. The pattern 1c shares some HEs with Indian patterns 1a and 1b, suggesting that also these cultivars derived from an ancestor originating in the Indian subcontinent.

#### Undescribed subgroup

The genotype ‘Chuoi mit’ (ITC1682) (misclassified as AB but determined to be triploid by Christelová *et al.* (2017)) exhibits its own specific HE pattern (pattern 7) (**Figure 4**). Morphological and genomic studies of additional genotypes would be required to support the creation of a new subgroup.

### West Oceania ABB

In this study, ABB accessions from Papua New Guinea (PNG) revealed a specific pattern (pattern 6). Given the absence in this analysis of genotypes belonging to the Kalapua subgroup, originating from PNG, we could not investigate its HE pattern. Therefore, it should be investigated whether Kalapua accessions share the same HE pattern 6 (**Figure 4**) or if an additional pattern from PNG exists. This would be a rationale to distinguish East ABB and West Oceania, usually combined, that however originated from different continents separated by the Wallace line.

### Genesis of HE patterns

The discovery of nine founding events opens the discussion on the possible routes that led to the creation of those different subgroups and formulate new hypotheses. To obtain a triploid genome constitution (2n=3x), a diploid gamete (n=2x) needs to meet a haploid gamete of the opposite sex (n=x). Usually, 2x gametes can be produced by tetraploid genotypes (2n=4x) or by a diploid genotype (2n=2x) through a meiosis failing one of the division (i.e. First or Second Division Restitution, FDR or SDR, respectively) and ending with a doubled chromosome set. Hybrids from different species are prone to set unreduced gametes, issued from FDR (Ramsey and Schemske 1998).

In banana, it has also been reported that allotriploid can produce viable reduced gametes (Shepherd 1999). To obtain an ABB constitution two gametic combinations (GCs) are consequently possible:

- AB x B: a 2x gamete containing A and B genomes (originating from i) a hybrid genotype (AB) by FDR, or ii) an allotriploid (AAB or ABB) or iii) an allotetraploid (AABB) that met a normal *M. balbisiana* gamete (B). In the first case the unreduced AB gamete brings 11 A and 11 B centromeres with possible HEs due to partial pairing between homoeologous chromosomes (**Figure 7**). In the second case HEs are also possible but an unbalanced A/B centromere representation is expected due to the double dose of one of the two subgenomes present in the allotriploid genotype. In the third case, 11 A and 11 B chromosomes are expected due to the regular meiosis between homologous chromosomes but, consequently, homoeologous pairing producing HEs is expected to be infrequent.
- A x BB: a haploid *M. acuminata* gamete (A) met a gamete (BB) originated from *M. balbisiana* by unreduced meiosis. Here, the recombination between A and B genome cannot take place before the allotriploidy establishment.

**Figure 7.**
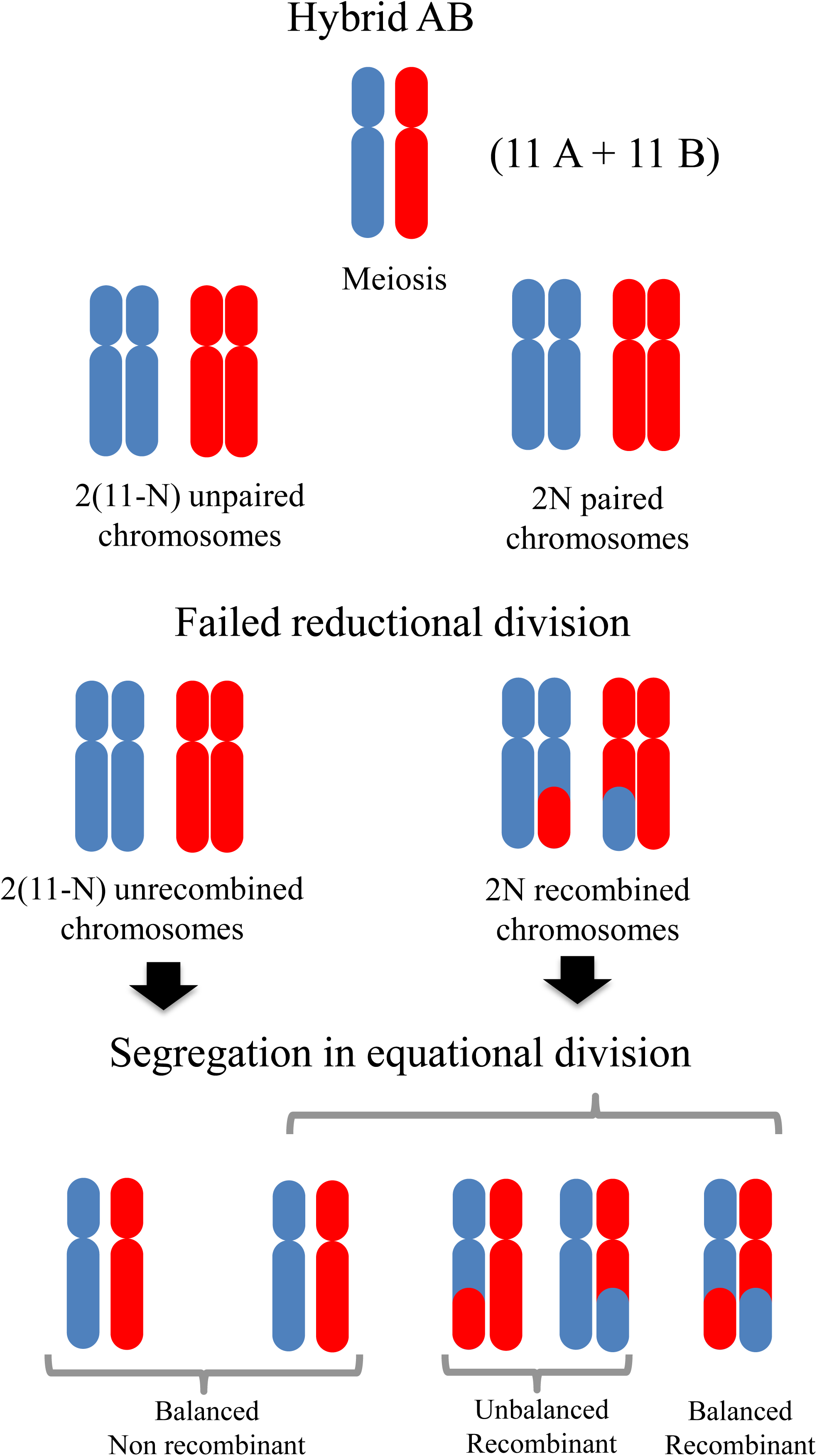
Scheme of backcrosses (BCs) generating introgressions of A genome in *M. balbisiana* parent of AB hybrids having genomic regions with unexpected subgenome ratio. In blue and red are represented chromosomes from *M. acuminata* (A genome) and *M. balbisiana* (B genome), respectively.

For the 2 genotypes from Indonesia without detected HEs, both GCs are possible and cannot be discriminated by our method due to the possible presence of invisible balanced HEs (see **Figure 7**). Such case of balanced HE was probably revealed in the second arm of chr05 thanks to the aneuploidy in ‘Simili Radjah’ genotype (**Figure 5d**) as none of the accessions with the same HE pattern (1b) showed deviating ratio for this chromosome (**Supplementary Material 2**).

#### Routes involving primary AB hybrids

Based on our observations of unbalanced regions between A and B genomes in allotriploids, we suggest that the diploid gamete contributing to the allotriploid with patterns 2, 3, 6 and 7 originated from a nucleus restitution during a diploid hybrid (AB) meiosis, which generated observed chromosome recombinations (**Figure 7**). De Langhe *et al.* 2010 hypothesized that this route would be restricted only to India due to the exclusive occurrence of AB in this country. However, since then, AB hybrids were also collected in Indonesia (Hermanto *et al.* 2014a; Hermanto *et al.* 2014b) making possible routes involving AB hybrids also for ABB genotypes with origin from South East Asia and New Guinea.

#### Routes leading to shared HEs

The observed HE patterns sharing some unbalanced regions **(Figure 2)** in HE patterns 1a, 1b and 1c cannot be explained with a simple AB x B gamete combination. In fact, identical or very similar deviating patterns (three unbalanced regions corresponding to these three recombination events) are unlikely to be obtained three times by fully independent meiosis. It must have been transmitted independently through common ancestry. In other words, the occurrence of partially shared patterns suggests repeated inheritance of the same HEs. Indeed, the reported differences in chloroplast origin (Carreel *et al.* 2002; Boonruangrod *et al.* 2008) support independent crossing pathways for patterns 1a, 1b and 1c (**Table 1**).

To explain this observation, we hypothesize that the shared HEs were already present in AB hybrids, who inherited them from gametes A and B previously introgressed. These hybrids then produced unreduced gametes in which new HEs possibly occurred during meiosis and gave birth to the allotriploid ancestors of the three patterns 1a, 1b and 1c. Some recombination events could also have even been provided by *M. balbisiana* haploid gametes contributing the second B subgenome. This route could be formulated as (A^b^B^a^) x B^a^B^a^ -> A^b^B^a^B^a^. A possible scenario driving to patterns 1a, 1b and 1c is proposed in **Supplementary Material 3**.

This hypothesis is supported by the observation of hybrid AB genotypes containing unbalanced regions (**Figure 1**) in which four different HE patterns were found. These results imply that the respective *M. balbisiana* donor plants were all introgressed with A genome portions. It suggests genetic fluxes between *M. acuminata* (or AA) and *M. balbisiana*, particularly in India, where these unbalanced AB hybrids and ABB allotriploid originated and where cultivated *M. balbisiana* are abundant (Subbaraya *et al.* 2006). Since in *Musa*, chloroplast and mitochondrion are inherited from the female and male gametes, respectively (Fauré *et al.* 1993), and both hybrid AB cytotypes have an A origin for both chloroplast and mitochondrion, the probable crossing scheme leading to the creation of these hybrids must have involved *M. acuminata* or AA gametes at least twice (**Figure 8**). However, it implies the incomplete sterility of AB hybrids that is able to produce not only unreduced gametes (n=2x=22) but also viable reduced gametes (n=x=11) where A and B chromosome recombine as in an intraspecific context and segregate.

**Figure 8.**
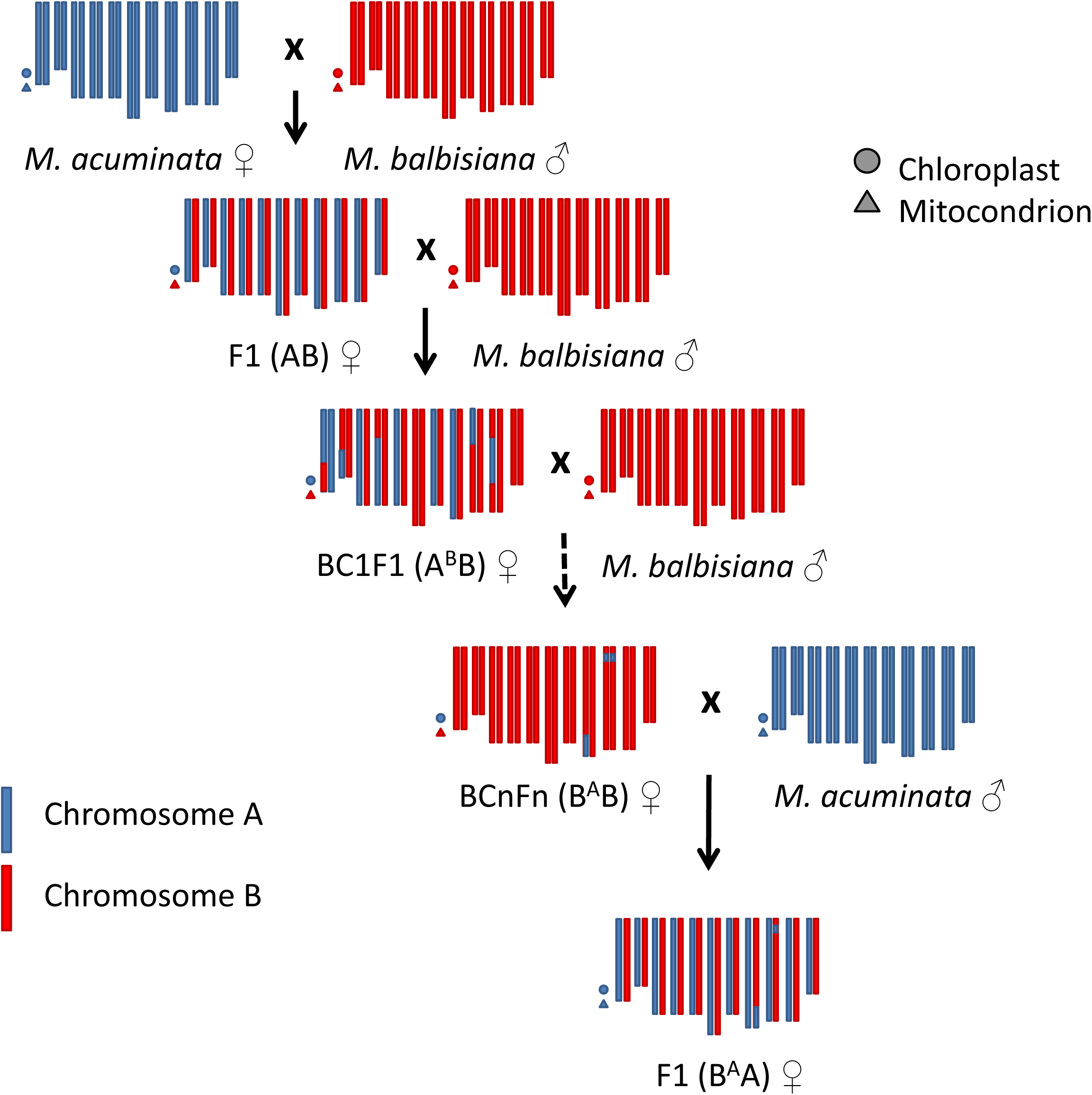
Scheme of hybrid AB meiosis producing unreduced gametes. In blue and red are represented chromosomes from *M. acuminata* (A genome) and *M. balbisiana* (B genome), respectively.

Interestingly in these genotypes, an interstitial region in chr09 first arm exhibits very similar HEs in allotriploids and diploids which again pleads for common ancestry (**Figure 3**). As already observed by Baurens *et al.* (2019), the subtelomeric region of chr09 first arm appears to be often enriched in the A genome (only Patterns 2, 3 and 2x-1 are not). In our sample, in addition to the above-mentioned diploid and triploid HE patterns, four patterns have independent HE where the A genome replaced the B counterpart in the subtelomeric region of chr09 first arm (**Figure 3**).

#### Routes leading to a B excess ratio

The patterns 4 and 5 are both characterized by a higher presence of the B genome, with three and six B centromeres having replaced the A counterpart, respectively. The absence of A centromeres (**Figure 4**) is not consistent with a meiosis of an AB hybrid where FDR gametes were produced by a failed first meiotic division. On the other hand, 2x gametes produced by the failing of equational division (SDR) are not expected to have an excess of the B genome or to miss centromeres with an A2:B1 constitution as observed in pattern 4 and 5.

The larger contribution of *M. balbisiana* to these cultivars is consistent with an unreduced n=2x gamete originating from a backcrossed diploid hybrid (originated by a n=x gamete from AB hybrid that met a n=x *M. balbisiana* gamete) crossed with a regular B gamete (**Figure 9**). An unreduced gamete from such backcrossed hybrid (A^b^B or B^a^B) is expected to have pairs of B centromeres for approximately half of its chromosomes and A/B heterozygous centromeres for the other half (**Figure 9**). It can be formulated as AB x BB -> A^b^B (or B^a^B) x BB -> A^b^BB (or B^a^BB).

**Figure 9.**
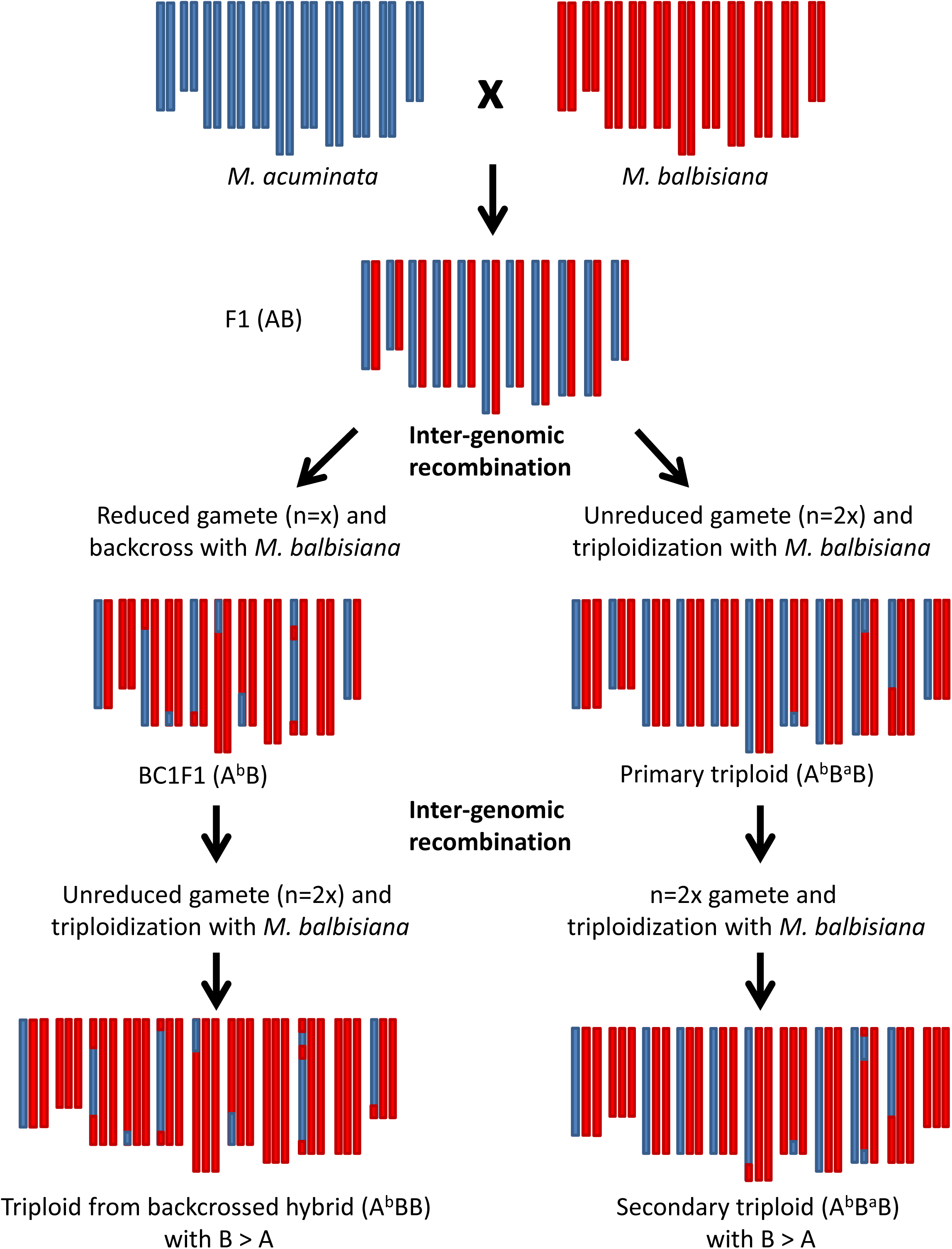
Scheme of possible crossing pathways generating B genome rich ABB patterns 4 and 5. In blue and red are represented chromosomes from *M. acuminata* (A genome) and *M. balbisiana* (B genome), respectively.

Alternatively, it can be hypothesized that a B-enriched n=2x gamete could be produced by an ABB allotriploid. In this case, according to random centromere segregation, one third of the centromeres are expected to be homozygous for B and two thirds are expected to be A/B heterozygous (**Figure 9**). It would be formulated as ABB x BB -> A^b^BB.

Pattern 4, with 3 on 11 centromeres having A0:B3 ratio, fits with the second hypothesis, whereas pattern 5, with 6 on 11 centromeres A0:B3, is more in agreement with first one. However, due to the low number of *Musa* chromosomes (11), those hypotheses cannot be excluded for both patterns. Somehow, independent origin of patterns 4 and 5 is supported by the completely different HEs and by the different crossing history that is inferred from the organelle origin. ‘Kluai Tiparot’ (pattern 5) has both organelles originating from *M. balbisiana*, whereas ‘Pelipita’ (pattern 4) has chloroplast inherited from *M. acuminata* and mitochondrion from *M. balbisiana* (Boonruangrod *et al.* 2008).

## Conclusions

The results of this study demonstrated multiple origins of ABB cultivars, with different and sometimes complex routes. It is possible that extension of RAD-Seq genome survey to other cultivars will allow the discovery of additional HE patterns, both in AB hybrids and in allotriploids. The frequent occurrences of HEs indicate that A and B subgenomes are prone to recombine, making the *M. balbisiana* genome a source of potential useful variability to create new cultivars able to answer the numerous challenges in banana breeding.

Compared with previous analysis methods to assess genetic diversity of the banana genepool, the whole genome survey based on NGS (RAD-Seq in this study) provides a detailed and exhaustive picture of the genome composition. Therefore, we recommend complementing SSR based characterization with NGS technologies such as RAD-Seq as a standard method to characterize the banana gene bank accessions and to classify them in an objective, precocious and repeatable way.

## Acknowledgements

We thank BGI for their technical assistance and services for the RAD sequencing. Our acknowledgements are also going to Matthew Turner from Amami Biodiversity Garden for permission of using *M. balbisiana* samples as reference. Finally, we thank Françoise Carreel, Angélique D’Hont and Franc-Christophe Baurens (CIRAD) for useful discussions. This work was supported by the Bioversity International project ‘Developing climate-smart bananas for the African Great Lakes region’, financed by the Belgian Development Cooperation and by donors through their contributions to the CGIAR Fund, and in particular to the CGIAR Research Program, Roots, Tubers and Bananas. The funders had no role in study design, data collection and analysis, interpretation of results, decision to publish, or preparation of the manuscript.

## Supplementary material

**Supplementary Material 1.** Number of SNPs assigned to each chromosome and to chloroplast/mitochondria (Cyto) in the analyzed cultivars.

**Supplementary Material 1**. Hidden HE revealed by aneuploidy. Chr05 pattern of “Blue Java” (euploid) and “Simili Radjah” (missing portion of second arm).

**Supplementary Material 2**. Schema of possible crossing pathways to HE pattern 1a, 1b and 1c. In green and red are represented chromosomes from *M. acuminata* (A genome) and *M. balbisiana* (B genome), respectively.

## Literature cited

Baurens F-C, Martin G, Hervouet C, et al. 2019. Recombination and Large Structural Variations Shape Interspecific Edible Bananas Genomes. Molecular Biology and Evolution 36: 97–111.

Boonruangrod R, Desai D, Fluch S, Berenyi M, Burg K. 2008. Identification of cytoplasmic ancestor gene-pools of Musa acuminata Colla and Musa balbisiana Colla and their hybrids by chloroplast and mitochondrial haplotyping. Theoretical and Applied Genetics 118: 43–55.

Carpentier SC, Panis B, Renaut J, et al. 2011. The use of 2D-electrophoresis and de novo sequencing to characterize inter- and intra-cultivar protein polymorphisms in an allopolyploid crop. Phytochemistry 72: 1243–1250.

Carreel F, de Leon DG, Lagoda P, et al. 2002. Ascertaining maternal and paternal lineage within Musa by chloroplast and mitochondrial DNA RFLP analyses. Genome 45: 679–692.

Cenci A, Hueber Y, Zorrilla-Fontanesi Y, et al. 2019. Effect of paleopolyploidy and allopolyploidy on gene expression in banana. BMC Genomics 20: 244.

Christelová P, Langhe ED, Hřibová E, et al. 2017. Molecular and cytological characterization of the global Musa germplasm collection provides insights into the treasure of banana diversity. Biodiversity and Conservation 26: 801–824.

Čížková J, Hřibová E, Humplíková L, Christelová P, Suchánková P, Doležel J. 2013. Molecular Analysis and Genomic Organization of Major DNA Satellites in Banana (Musa spp.). PLoS ONE 8: e54808.

Daniells J. 2001. Musalogue: a catalogue of Musa germplasm: diversity in the genus Musa. Bioversity International.

De Langhe E. 2002. Banana Diversity in the Middle East (Jordan, Egypt and Oman). International Network for the Improvement of Banana and Plantain (INIBAP), Montpellier, France: 1–56.

De Langhe E, Hřibová E, Carpentier S, Doležel J, Swennen R. 2010. Did backcrossing contribute to the origin of hybrid edible bananas? Annals of Botany 106: 849–857.

De Langhe E, Langhe ED, Vrydaghs L, Maret P de, Perrier X, Denham T. 2009. Why Bananas Matter: An introduction to the history of banana domestication. Ethnobotany Research and Applications 7: 165–177.

D’Hont A, Denoeud F, Aury J-M, et al. 2012. The banana (Musa acuminata) genome and the evolution of monocotyledonous plants. Nature 488: 213.

D’Hont A, Paget-Goy A, Escoute J, Carreel F. 2000. The interspecific genome structure of cultivated banana, Musa spp. revealed by genomic DNA in situ hybridization. Theoretical and Applied Genetics 100: 177–183.

Droc G, Lariviere D, Guignon V, et al. 2013. The Banana Genome Hub. Database 2013: bat035–bat035.

Fauré S, Noyer JL, Horry JP, Bakry F, Lanaud C, Gonzalez de León D. 1993. A molecular marker-based linkage map of diploid bananas (Musa acuminata). Theoretical and Applied Genetics 87: 517–526.

Hermanto C, Sutanto A, Daniells J, Riska, HS E, Alfons, et al. 2014. Triangle Banana Exploration Report, North Sulawesi and North Maluku, Indonesia. 6-24 October 2012. Bioversity International.

Hermanto C, Sutanto A, Daniells J, Riska, HS E, Hilman Y, et al. 2014. Triangle Banana Exploration Report, Central Maluku and Lesser Sunda Islands, Indonesia. 16 February - 6 March 2013. Bioversity International.

Hippolyte I, Jenny C, Gardes L, et al. 2012. Foundation characteristics of edible Musa triploids revealed from allelic distribution of SSR markers. Annals of Botany.

Hřibová E, Čížková J, Christelová P, Taudien S, Langhe E de, Doležel J. 2011. The ITS1-5.8S-ITS2 Sequence Region in the Musaceae: Structure, Diversity and Use in Molecular Phylogeny. PLOS ONE 6: e17863.

Janssens SB, Vandelook F, De Langhe E, et al. 2016. Evolutionary dynamics and biogeography of Musaceae reveal a correlation between the diversification of the banana family and the geological and climatic history of Southeast Asia. The New Phytologist 210: 1453–1465.

Jenny C, Carreel F, Bakry F. 1997. Revision on banana taxonomy: Klue Tiparot (Musa sp) reclassified as a triploid. Fruits 2: 83–91.

Jenny C, Holtz Y, Horry J-P, Bakry F. 2011. Synthesis of new interspecific triploid hybrids from natural AB germplasm in banana (Musa sp.) In: VII International Symposium on Banana: ISHS-ProMusa Symposium on Bananas and Plantains: Towards Sustainable Global Production 986. 209–217.

Jeridi M, Bakry F, Escoute J, et al. 2011. Homoeologous chromosome pairing between the A and B genomes of Musa spp. revealed by genomic in situ hybridization. Annals of Botany.

Karamura E, Frison E, Karamura D, Sharrock S. 1998. Banana production systems in eastern and southern Africa. Bananas and food security. INIBAP, Montpellier: 401–412.

Kissel E, van Asten P, Swennen R, Lorenzen J, Carpentier SC. 2015. Transpiration efficiency versus growth: Exploring the banana biodiversity for drought tolerance. Scientia Horticulturae 185: 175–182.

Li H, Durbin R. 2009. Fast and accurate short read alignment with Burrows–Wheeler transform. Bioinformatics 25: 1754–1760.

Martin M. 2011. Cutadapt removes adapter sequences from high-throughput sequencing reads. EMBnet.journal 17: 10.

Martin G, Baurens F-C, Droc G, et al. 2016. Improvement of the banana “Musa acuminata” reference sequence using NGS data and semi-automated bioinformatics methods. BMC genomics 17: 243.

McKenna A, Hanna M, Banks E, et al. 2010. The Genome Analysis Toolkit: a MapReduce framework for analyzing next-generation DNA sequencing data. Genome Research 20: 1297–1303.

Noumbissié GB, Chabannes M, Bakry F, et al. 2016. Chromosome segregation in an allotetraploid banana hybrid (AAAB) suggests a translocation between the A and B genomes and results in eBSV-free offsprings. Molecular Breeding 36: 38.

Perrier X, Langhe ED, Donohue M, et al. 2011. Multidisciplinary perspectives on banana (Musa spp.) domestication. Proceedings of the National Academy of Sciences 108: 11311–11318.

Ramsey J, Schemske DW. 1998. Pathways, mechanisms, and rates of polyploid formation in flowering plants. Annual Review of Ecology and Systematics 29: 467–501.

Ravi I, Uma S, Vaganan MM, Mustaffa MM. 2013. Phenotyping bananas for drought resistance. Frontiers in Physiology 4.

Rosales F, Arnaud E, Coto J. 1999. A tribute to the work of Paul Allen. A catalogue of wild and cultivated bananas. INIBAP, Montpellier, France.

Ruas M, Guignon V, Sempere G, et al. 2017. MGIS: managing banana (Musa spp.) genetic resources information and high-throughput genotyping data. Database: The Journal of Biological Databases and Curation 2017.

Sales EK, Butardo NG, Paniaga HG, Jansen H, Dolezel J. 2010. Assessment of ploidy and genome constitution of some Musa balbisiana cultivars using DArT markers. Philippine Journal of Crop Science (Philippines).

Saraswathi MS, Uma S, Vadivel E, et al. 2011. Diversity analysis in Indian cooking bananas (Musa, ABB) through morphotaxonomic and molecular characterisation. Acta Horticulturae: 123–131.

Sardos J, Perrier X, Doležel J, et al. 2016. DArT whole genome profiling provides insights on the evolution and taxonomy of edible Banana (Musa spp.). Annals of Botany: mcw170.

Sempéré G, Philippe F, Dereeper A, Ruiz M, Sarah G, Larmande P. 2016. Gigwa—Genotype investigator for genome-wide analyses. GigaScience 5.

Shepherd K. 1999. Cytogenetics of the genus Musa. IPGRI.

Simmonds NW. 1962. The evolution of the bananas. London (GBR): Longmans.

Simmonds NW, Shepherd K. 1955. The taxonomy and origins of the cultivated bananas. Journal of the Linnean Society of London, Botany 55: 302–312.

Subbaraya U, Lutaladio N, Baudoin WO, others. 2006. Farmer’s knowledge of wild Musa in India. Plant Production and Protection Division, FAO, Rome: 1–46.

Thomas DS, Turner DW, Eamus D. 1998. Independent effects of the environment on the leaf gas exchange of three banana (Musa sp.) cultivars of different genomic constitution. Scientia Horticulturae 75: 41–57.

Valmayor R, Jamaluddin S, Silayoi B, et al. 1999. Banana cultivar names and synonyms in Southeast Asia. Advancing Banana and Plantain R & D in Asia and the Pacific.

Valmayor R, Silayoi B, Jamaluddin S, Kusomo S, Espino R, Pascua O. 1991. Banana classification and commercial cultivars in Southeast Asia.

Vanhove A, Garcia S, Swennen R, Panis B, Carpentier SC. 2012. Understanding Musa drought stress physiology using an autotrophic growth system. Communications in agricultural and applied biological sciences 77: 89–93.

van Wesemael J, Kissel E, Eyland D, Lawson T, Swennen R, Carpentier S. 2019. Using Growth and Transpiration Phenotyping Under Controlled Conditions to Select Water Efficient Banana Genotypes. Frontiers in Plant Science 10.

