## Supplementary Material 1 for "Unraveling the complex story of intergenomic recombination in ABB allotriploid bananas"

| Accession code | Accession name | Genome | A assigned | B assigned | Chr01 | Chr02 |
|---|---|---|---|---|---|---|
| ITC0026 | Sabra | ABB | 124937 | 95130 | 16769 | 16328 |
| ITC0053 | Bom | ABB | 112499 | 93372 | 15185 | 15755 |
| ITC0087 | Kayinja | ABB | 70124 | 63202 | 9868 | 9836 |
| ITC0101 | Fougamou 1 | ABB | 85157 | 81820 | 12377 | 12528 |
| ITC0123 | Simili Radjah | ABB | 94378 | 126981 | 16691 | 16310 |
| ITC0245 | Safet Velchi | AB | 85032 | 146993 | 16194 | 16749 |
| ITC0339 | Pisang Awak | ABB | 80026 | 83764 | 12197 | 12275 |
| ITC0361 | Blue Java | ABB | 91203 | 91188 | 13627 | 13236 |
| ITC0396 | Pelipita | ABB | 70196 | 90890 | 13069 | 6928 |
| ITC0397 | Pelipita Majoncho | ABB | 64657 | 85323 | 12113 | 6404 |
| ITC0472 | Pelipita | ABB | 91973 | 72400 | 13291 | 6937 |
| ITC0473 | Balonkawe | ABB | 38368 | 72350 | 11465 | 5596 |
| ITC0526 | Kluai Namwa Khom | ABB | 87117 | 83644 | 12697 | 12701 |
| ITC0632 | Cachaco enano | ABB | 103950 | 92911 | 14746 | 14501 |
| ITC0643 | Cachaco | ABB | 83587 | 86443 | 12866 | 12339 |
| ITC0652 | Kluai Tiparot | ABB | 50295 | 99288 | 15707 | 7429 |
| ITC0659 | Namwa Khom | ABB | 84679 | 87552 | 12744 | 12858 |
| ITC0767 | Dole | ABB | 85294 | 93465 | 14111 | 13876 |
| ITC0983 | Auko | ABB | 92804 | 101886 | 14278 | 14273 |
| ITC0987 | Auko | ABB | 92994 | 105912 | 14624 | 14548 |
| ITC0990 | Vunapope | ABB | 98191 | 147374 | 18231 | 18092 |
| ITC1034 | Kunnan | AB | 153665 | 87459 | 17673 | 17507 |
| ITC1138 | Saba | ABB | 79331 | 88841 | 12938 | 9304 |
| ITC1483 | Monthan | ABB | 53555 | 49010 | 7794 | 7630 |
| ITC1599 | Kambani | ABB | 87857 | 84808 | 12797 | 12977 |
| ITC1600 | INIVIT PB-2003 | ABB | 90945 | 85362 | 13519 | 9646 |
| ITC1682 | Chuoi mit | ABB | 86553 | 89061 | 10112 | 12751 |
| ITC1700 | Kepok Kuning | ABB | 96150 | 92011 | 13557 | 13390 |
| ITC1719 | Chinia | ABB | 98289 | 90727 | 13800 | 13913 |
| ITC1721 | Karpuravalli | ABB | 91978 | 91095 | 13609 | 13779 |
| ITC1728 | Sambrani Monthan | ABB | 81924 | 86054 | 12565 | 12429 |
| ITC1729 | Padali Moongil | AB | 125662 | 78178 | 14805 | 14568 |
| ITC1737 | Ya Khine | ABB | 62811 | 78051 | 10538 | 11123 |
| ITC1738 | Kyauk Sein Phee Kyan | ABB | 99803 | 69010 | 12897 | 12998 |
| ITC1745 | Kepok Tanjung | ABB | 94311 | 91072 | 13514 | 13309 |
| ITC1746 | Bankel | ABB | 91356 | 91136 | 13793 | 13432 |
| ITC1747 | Agniswar | AB | 98785 | 63715 | 11940 | 11526 |
| ITC1748 | Boddida Bukkisa | ABB | 85098 | 81985 | 12631 | 12182 |
| ITC1749 | Vananthpurani | ABB | 90066 | 99489 | 13778 | 13878 |
| ITC1750 | Ney Vannan | ABB | 85792 | 87206 | 13066 | 12623 |
| ITC1751 | Adukka Kunnan | AB | 80092 | 53897 | 9995 | 9633 |
| ITC1752 | Poovilla Chundan | AB | 134680 | 79919 | 15656 | 15392 |
| Sum002 | Muku Bugis | AB | 173635 | 97945 | 19631 | 19582 |
| Sum004 | Mu'u Seribu | AB | 166545 | 95682 | 19296 | 19196 |
| Sup008 | Mu’u Pundi | AB | 173702 | 97163 | 19664 | 19457 |

| Chr03 | Chr04 | Chr05 | Chr06 | Chr07 | Chr08 | Chr09 | Chr10 | Chr11 | Total | Cyto |
|---|---|---|---|---|---|---|---|---|---|---|
| 22430 | 22393 | 20786 | 25483 | 19147 | 23199 | 20829 | 20492 | 9976 | 217832 | 2235 |
| 21304 | 21406 | 20061 | 23688 | 17815 | 21452 | 19446 | 12261 | 15399 | 203772 | 2099 |
| 13211 | 14427 | 12607 | 15576 | 11036 | 13691 | 12785 | 8969 | 9927 | 131933 | 1393 |
| 16928 | 17711 | 16087 | 19502 | 13841 | 17503 | 15942 | 10580 | 12215 | 165214 | 1763 |
| 22355 | 22521 | 20200 | 26176 | 19480 | 23265 | 20772 | 20596 | 10833 | 219199 | 2160 |
| 23277 | 26844 | 21696 | 24803 | 20033 | 23229 | 21160 | 19698 | 16025 | 229708 | 2317 |
| 16603 | 17352 | 15917 | 19113 | 13418 | 17248 | 15644 | 10154 | 12065 | 161986 | 1804 |
| 18408 | 18994 | 17290 | 21174 | 15474 | 18726 | 17267 | 16779 | 9477 | 180452 | 1939 |
| 17669 | 21407 | 16555 | 19437 | 9029 | 17846 | 16848 | 13894 | 6735 | 159417 | 1669 |
| 16331 | 19999 | 15332 | 18111 | 8526 | 16672 | 15684 | 12804 | 6451 | 148427 | 1553 |
| 18098 | 21959 | 16902 | 19873 | 9120 | 18326 | 17158 | 14199 | 6831 | 162694 | 1679 |
| 11287 | 10224 | 14906 | 10918 | 8344 | 7478 | 12888 | 6896 | 10716 | 110718 | 0 |
| 17311 | 18001 | 16424 | 20019 | 14176 | 17779 | 16177 | 11018 | 12651 | 168954 | 1807 |
| 19837 | 20375 | 18716 | 23203 | 16787 | 20459 | 18796 | 18163 | 9236 | 194819 | 2042 |
| 17132 | 17774 | 15952 | 19982 | 14314 | 17834 | 16077 | 15667 | 8324 | 168261 | 1769 |
| 14990 | 13743 | 19487 | 14847 | 11439 | 10167 | 16697 | 9310 | 14320 | 148136 | 1447 |
| 17540 | 18320 | 16438 | 20070 | 14426 | 18038 | 16367 | 10962 | 12648 | 170411 | 1820 |
| 19086 | 19599 | 17785 | 22172 | 16050 | 9585 | 17953 | 17618 | 8957 | 176792 | 1967 |
| 17827 | 18497 | 18686 | 22179 | 16489 | 19900 | 18506 | 18158 | 13846 | 192639 | 2051 |
| 18277 | 18936 | 19297 | 22489 | 17122 | 20222 | 18889 | 18072 | 14247 | 196723 | 2183 |
| 22166 | 22334 | 23711 | 28360 | 21200 | 25126 | 23328 | 22754 | 17824 | 243126 | 2439 |
| 23900 | 28862 | 23135 | 26321 | 19825 | 23074 | 21006 | 20921 | 16544 | 238768 | 2356 |
| 17394 | 17187 | 16224 | 20313 | 14548 | 18034 | 16078 | 15918 | 8427 | 166365 | 1807 |
| 10293 | 11063 | 9681 | 11684 | 8616 | 10046 | 9719 | 9645 | 5325 | 101496 | 1069 |
| 17569 | 18372 | 16600 | 20156 | 14225 | 18065 | 16343 | 11061 | 12641 | 170806 | 1859 |
| 18459 | 17787 | 17168 | 21062 | 15329 | 18779 | 16829 | 17133 | 8740 | 174451 | 1856 |
| 17614 | 18338 | 16695 | 20356 | 14589 | 18186 | 16564 | 15863 | 12677 | 173745 | 1869 |
| 17972 | 21784 | 17058 | 21301 | 15542 | 18640 | 17096 | 16703 | 13240 | 186283 | 1878 |
| 18809 | 21612 | 17237 | 21524 | 16286 | 19371 | 17625 | 13187 | 13650 | 187014 | 2002 |
| 18664 | 19233 | 17716 | 21330 | 15303 | 19125 | 17387 | 11665 | 13304 | 181115 | 1958 |
| 17000 | 17682 | 16066 | 19667 | 14231 | 16668 | 15998 | 15578 | 8319 | 166203 | 1775 |
| 20125 | 23662 | 19188 | 22354 | 17013 | 19736 | 18111 | 17520 | 14576 | 201658 | 2182 |
| 14401 | 14653 | 14061 | 16262 | 12135 | 14788 | 13807 | 8418 | 10676 | 140862 | 0 |
| 17210 | 17199 | 16271 | 19972 | 14473 | 16896 | 16682 | 15938 | 8277 | 168813 | 0 |
| 17760 | 21350 | 16771 | 20881 | 15198 | 18532 | 16713 | 16670 | 12860 | 183558 | 1825 |
| 18430 | 18902 | 17309 | 21196 | 15450 | 19011 | 17380 | 16873 | 8743 | 180519 | 1973 |
| 15711 | 19323 | 15354 | 18094 | 13502 | 15267 | 14383 | 13881 | 11688 | 160669 | 1831 |
| 16798 | 17492 | 15867 | 19460 | 14211 | 17220 | 15943 | 15349 | 8160 | 165313 | 1770 |
| 18979 | 21545 | 17450 | 21554 | 16450 | 19411 | 17672 | 13150 | 13703 | 187570 | 1985 |
| 17323 | 17825 | 16398 | 20149 | 14695 | 17928 | 16360 | 15716 | 9077 | 171160 | 1838 |
| 12935 | 15927 | 12373 | 14950 | 10707 | 13008 | 11920 | 11323 | 9668 | 132439 | 1550 |
| 21418 | 24913 | 20266 | 23241 | 17882 | 20737 | 19289 | 18317 | 15311 | 212422 | 2177 |
| 26546 | 31638 | 24682 | 30178 | 22827 | 26566 | 24571 | 23885 | 18783 | 268889 | 2691 |
| 25073 | 29340 | 23509 | 29687 | 22153 | 25967 | 23369 | 23236 | 18707 | 259533 | 2694 |
| 26406 | 31541 | 24382 | 30214 | 22786 | 26596 | 24481 | 23726 | 18827 | 268080 | 2785 |
