## Supplementary figures and images for "Unraveling the complex story of intergenomic recombination in ABB allotriploid bananas"

### Supplementary Material 2

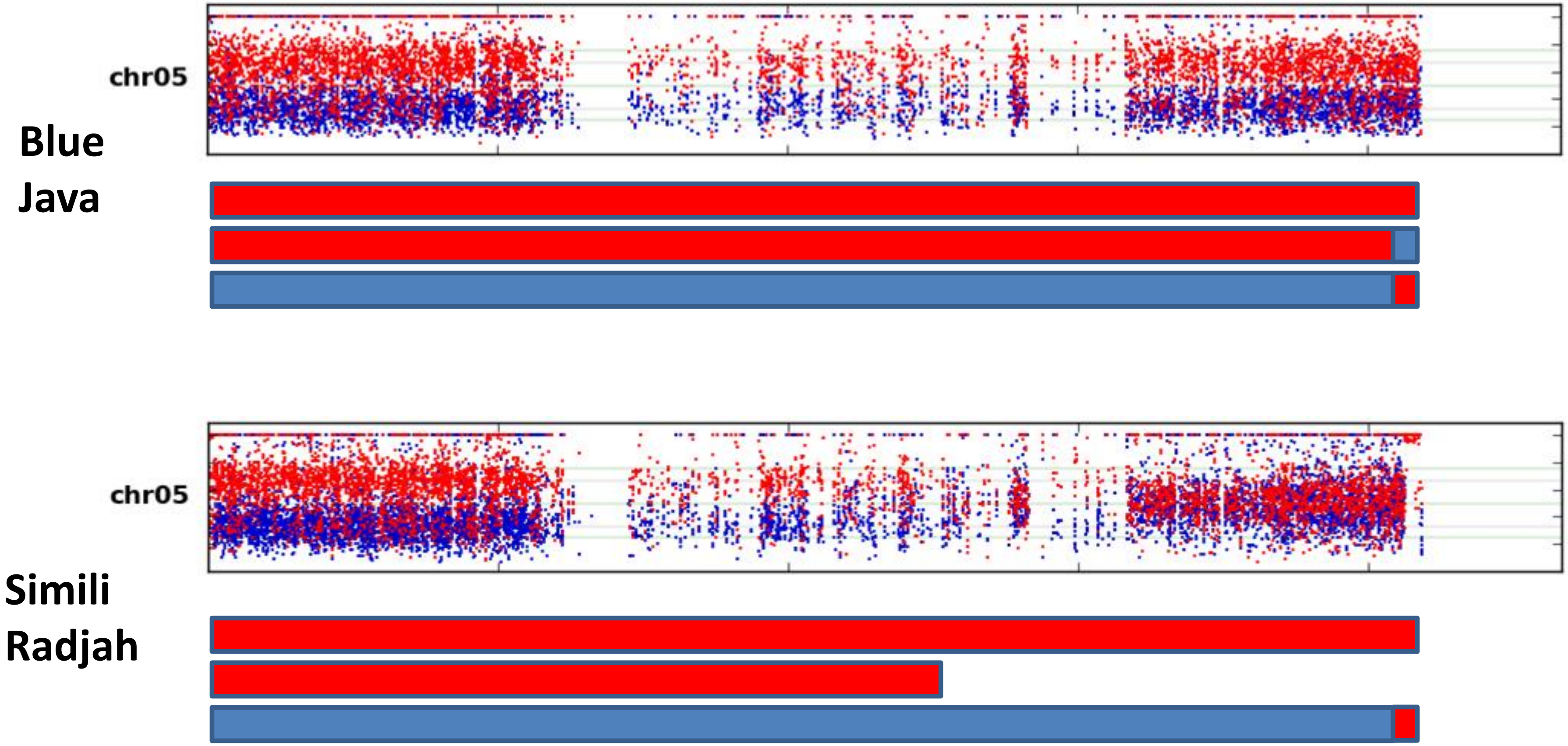

### Supplementary Material 3

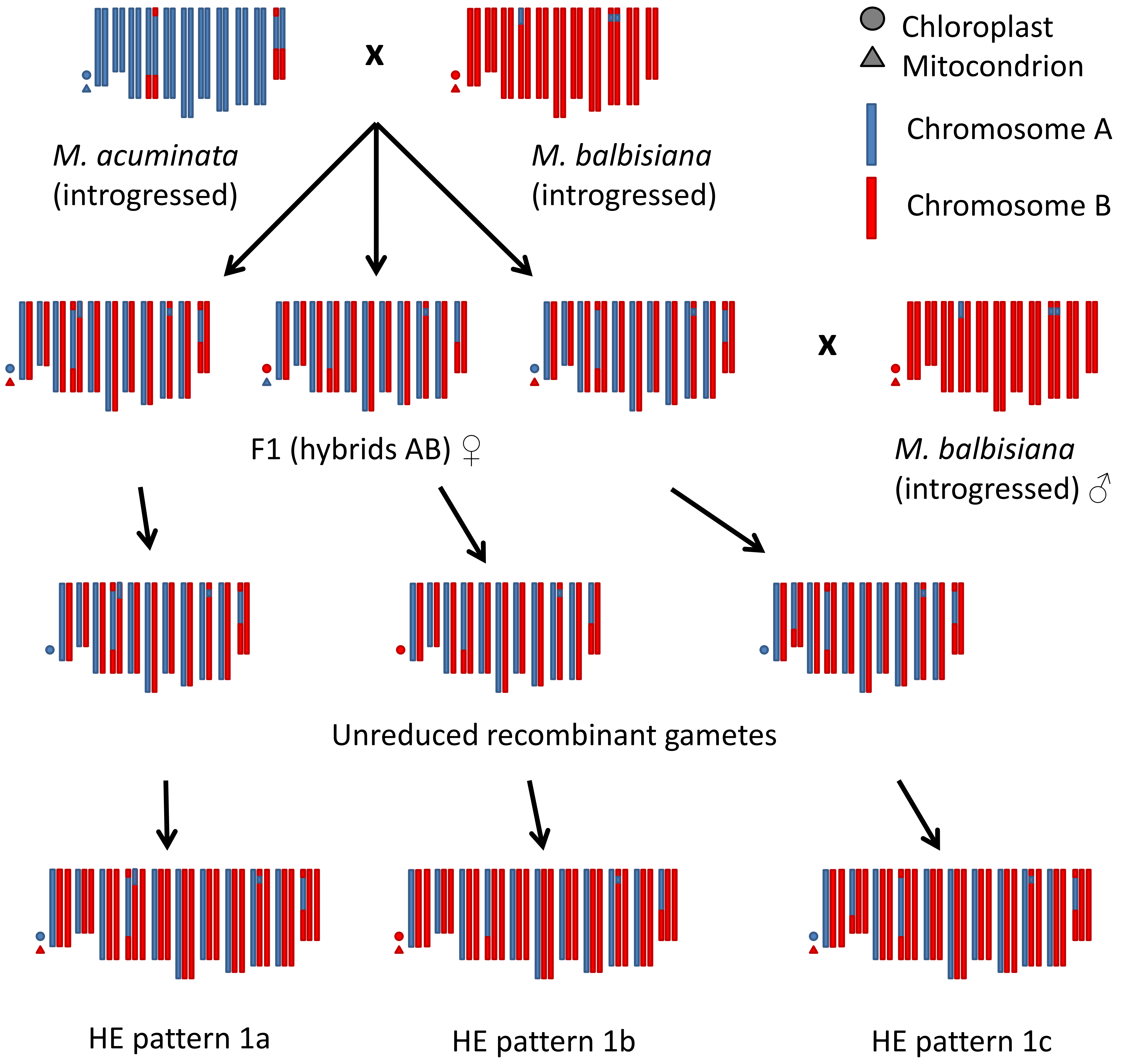
